# Short oligomers rather than rings of human RAD52 promote single-strand annealing

**DOI:** 10.1101/2023.08.11.553006

**Authors:** Maria A. Kharlamova, Manish S. Kushwah, Tobias J. Jachowski, Sivaraman Subramaniam, A. Francis Stewart, Philipp Kukura, Erik Schäffer

## Abstract

Genome maintenance and stability rely on the repair of DNA double-strand breaks. The break repair can be mediated by the single-strand annealing protein RAD52. RAD52 forms rings that are thought to promote annealing. However, RAD52’s annealing activity decreases with increasing concentrations that favor ring formation. Thus, which oligomeric form and how RAD52 anneals DNA strands and detects sequence homology is unclear. We combine mass photometry with biochemical assays to quantify oligomeric states of human RAD52 with and without DNA and put forward an alternative mechanism illustrating the critical role of short oligomers for single-stranded DNA annealing. We found that while truncated RAD52 formed undecameric rings at nanomolar concentrations, full-length RAD52 was mostly monomeric at lower nanomolar, physiological concentrations. At higher concentrations, it formed rings with a variable stoichiometry from heptamers to tridecamers. At low concentrations, with hardly any rings present, RAD52 already promoted single-strand annealing. Rings and short oligomers could bind at least two single DNA strands, but if complementary strands were both bound to rings annealing was inhibited. Our findings suggest that single-strand annealing and homology detection is mediated by short oligomers of RAD52 instead of rings.

## Introduction

RAD52 is a DNA single-strand annealing protein that is highly conserved in eukaryotes. Unlike RAD51/RecA, RAD52 promotes double strand break repair without utilizing ATP^1^. RAD52 also contributes to D-loop formation^2^, releases stalled replication forks during break-induced replication^3,4^ and is involved in double strand break repair at G-quadruplex-forming sequences^5^. In yeast, Rad52 is essential for homologous recombination^6,7^ by interacting with replication protein A (RPA) coated single-stranded DNA (ssDNA) to mediate the loading of Rad51^8,9^. In mammals, this mediator activity is performed by BRCA2 that outcompetes RAD52^10^. In BRCA2-deficient cancer cells, RAD52 takes over^11^ and, depending on RPA’s phosphorylation state^12^, either acts as a mediator for homologous recombination or promotes single strand annealing making it a potential therapeutic target for cancer treatment^13,14^. To utilize RAD52 as a cancer target and understand its various functions, it is essential to understand the primary function of RAD52 as a single-strand annealing protein.

C-terminally truncated RAD52, typically at amino acid 209 or 212, is sufficient for single-strand annealing and oligomerization^15–17^. This domain self-associates to form rings that can self-associate further to form larger aggregates^18,19^. Early electron microscopy images described RAD52 as a heptameric ring (7mers) with long protrusions attributed to the C-terminal domain^19,20^. Further work showed that human RAD52 can also form undecameric rings (11mers)^21,22^ while yeast Rad52 forms decameric rings (10mers)^23^. Undecamers are also formed by the C-terminal truncation RAD52^24–26^. The rings have two DNA-binding sites^2^. One binding site, located within the inner groove of the ring, is specific to ssDNA and important for annealing^2,27^. At this site, four nucleotides are bound per RAD52 monomer^16,28^. The second DNA binding site is located on the ring’s outer part near the groove and can also bind double-stranded DNA (dsDNA), the latter being important for D-loop formation^2,29^. Furthermore, this site promotes aggregation of RAD52 in the presence of ssDNA^27^.

Apart from the conspicuous ring structures, a range of shorter oligomers and large aggregates have also been observed^19^. Based on drug interactions, RAD52 dimers appear to be a stable oligomeric form^30^. Also, RAD52 rings and aggregates can be disassembled by ssDNA, magnesium ions or RPA^21,31^. In ssDNA-RPA-RAD52 complexes, rings were not present suggesting that the active form of RAD52 in this complex is monomeric^21^. RAD52 binds stably to ssDNA and in a diffusive manner to dsDNA^32^. However, the oligomeric state of the bound molecules could not be clearly identified. Thus, it remains unclear whether short oligomers bind DNA equally well as the rings.

Despite the evidence that RAD52 multimerization is dynamic and variable, current DNA annealing models are based on rings^2,16,27,33–35^. According to a recent model, two ssDNA overhangs formed after a double-strand break are bound to the inner groove of two RAD52 rings. Rings slide or rotate along each other to identify homology. Once the homology is detected, ssDNA molecules relocate from the inner grooves to the outer binding sites where annealing is completed. Notably the authors acknowledge that the distance between ring-bound ssDNAs is too large for homology detection and additional mechanisms or conformational changes are necessary^27^. For a homology search, eight contiguous bases are proposed to be efficient, while for unique homology detection, 12 to 17 contiguous bases appear to be required^36–39^. Therefore, for homology detection, at least three to four consecutive protomers—the subunits of an oligomeric protein—from one ring, each binding four bases^16,28^, should be able to interact simultaneously with the corresponding number of protomers from a neighboring ring. This simultaneous interaction is unlikely because of the ring curvature. To alleviate the conceptual problems, partial peeling off of at least one of the strands has been suggested^35^. While structures of truncated RAD52 with ssDNA were solved^27^, no structure of RAD52 with annealed DNA has been solved so far^26^.

Using DNA annealing assays, several prokaryotic single-strand annealing proteins have been biochemically identified. These proteins also multimerize and, in the case of lambda phage Red*β*, form un- and dodecadecameric rings (11,12mers)^40^, classification by sequence homology led to the identification of three separate single-strand annealing protein classes named after their most prominent members; RAD52, Red*β* /RecT and Erf^41^. Using advanced bioinformatics, a very distant short tripartite signature shared between RAD52 and the Red*β* /RecT class was identified^42^. The deep relationship between the three single-strand annealing protein classes was secured recently when the first Red*β* /RecT structures were obtained by cryogenic electron microscopy (cryo-EM)^43,44^. Both structures were clearly similar to the known structure of the RAD52 annealing domain thereby revealing a new protein fold that anchors the RAD52 single-strand annealing protein superfamily, which also includes Erf^26^. Both cryo-EM structures are multimeric single-strand-annealing-protein– annealed-DNA filaments that cast light on the annealing mechanism and lend credence to the monomer to multimer model that has arisen from studies with Red*β* ^37, 42, 45–47^.

The RAD52 C-terminal domain is intrinsically disordered and contains RPA and RAD51 binding sites that by and large do not affect RAD52’s annealing activity^9,22,27^. Furthermore, it contains a binding site for self-association of rings^19^ that is inhibited by RPA^48^. The C-terminal domain also contains a weak nuclear localization sequence that requires RAD52 oligomerization for nuclear localization^49^. Notably, interaction of E. coli SSB, the prokaryotic counterpart of RPA, with the C-terminal domain of Red*β* has recently been identified, thereby adding further functional weight to the RAD52 single-strand annealing protein superfamily proposition^47,50^.

RAD52’s annealing activity has been thoroughly studied^15,16^. With increasing RAD52 concentration, annealing rates and product yield first increase and then decrease again^9,35,51,52^ with the maximum occurring in the lower nanomolar range. This maximum roughly matches the estimated *in vivo* concentration of 20 nM in yeast and 2 nM in humans^53–55^. Furthermore, efficient annealing of human RAD52 has been reported down to 1 nM^52^.

For the annealing reaction, the order of incubation of the catalyst RAD52 with its two ssDNA substrates is unclear. Usually, one strand is incubated with the protein for some time and then the other strand is added to the reaction^51,52^. When incubating both complementary strands separately with the protein before mixing them, one study reported a 2.5-fold decrease in annealing efficiency^51^, while other experiments required such an incubation^33^. In general, RAD52 ring structures have been observed at high concentrations, i.e., at conditions that inhibit annealing. No study has so far directly visualized the nucleoprotein complex during annealing^26^. Thus, while RAD52 anneals DNA, the oligomeric state that promotes annealing and how homology is detected are still unclear.

To shed more light on how RAD52 anneals DNA, we characterized the oligomeric state of full-length, human RAD52 and a truncated version at nanomolar concentrations by using mass photometry^57^. We complemented the experiments with electromobility shift assays and compared how the oligomeric states of both proteins are altered by addition of complementary ssDNA. Our findings agree with a dynamic annealing mechanism that employs RAD52 short oligomers.

## Materials and Methods

### Single-molecule mass photometry

Mass photometry was performed on a commercial instrument OneMP (REFEYN Inc., Oxford, United Kingdom) using the regular field of view for the measurements and standard averaging parameters. Only for the RAD52 annealing measurements (Fig. 7), the medium field of view was used to increase RAD52 counts. The mass range of single, label-free proteins in their native state that can be detected quantitatively is 40 kDa–5 MDa. Samples were diluted in Buffer 1 (Tris-HCl 20 mM, NaCl 50 mM, pH 7.5). Purified Type 1 water (18.2 MΩ cm, Nanopure System MilliQ reference with Q-POD and Biopak filter) was used for all experiments. Buffers were filtered two times (0.22 *μ*m pore size filter units, Merck) and degassed for 30 min. Glass slides (25 *×* 50 mm) were sonicated first in water, then in isopropanol, and again in water, each time 5 min at room temperature. Afterward slides were dried with filtered air. Cell culture gaskets were placed on top of the slides to contain the samples. The system was calibrated with NativeMark Unstained Protein Standard (Invitrogen, Cat. #LC0725, 1:400 dilution). Buffer 1 was measured on each day prior to experiments. Acquired movies with binding events were processed in DiscoverMP (REFEYN Inc.). Generated HDF5 files were processed and fitted using a custom-written Python script. Resulting fits represent envelopes of multi-Gaussian fits for each oligomeric state of the protein. Peaks with areas below 1 % were excluded from the analysis. The standard deviations were fixed to the value of the standard deviation of the calibration proteins. The best fit was chosen based on the Akaike criterium.

### Protein purification

The pET-15b vector harboring full length human RAD52 and truncated RAD52(209) expression plasmids were gifts from Prof. Wataru Kagawa (Meisei University, Tokyo, Japan) and Prof. S.C. West (Cancer Research UK), respectively. To overexpress proteins, the *E. coli* strain BL21(DE3) was cotransformed with the pET-15b vector containing the RAD52 genes and the pRARE vector, encoding low abundance tRNAs. For the purification, 6 and 4 liter for RAD52 and RAD52(209), respectively, of a lysogeny broth culture supplemented with ampicillin and chloramphenicol were inoculated with fresh overnight culture and incubated at 37 °C, and protein expression was induced at an optical density (A_600) of 0.8 with 0.5 mM isopropyl 1-thio-*β* -D-galactopyranoside (final concentration). Subsequently, the temperature was lowered to 24 °C. Induction was carried out overnight. After induction, cells were harvested, resuspended in ≈ 80 ml of Buffer A (50 mM Tris-HCl, pH 7.8, 1.0 M KCl, 2 mM 2-mercaptoethanol, and 10 % glycerol) containing 10 mM imidazole and a protease inhibitor cocktail (Merck KgaA, Germany), and lysed by sonication. All procedures after cell harvesting were performed at 4 °C. The cell lysate was cleared of insoluble material by centrifugation at 40,000 g for 60 min. The supernatant was loaded onto a Buffer-A-equilibrated His-trap Ni-NTA fastflow (5 ml) column (Cytiva) used with an ÄKTAexplorer (Cytiva). After loading cell lysates, washing was carried out for 25 column volumes (125 ml) using Buffer A. Proteins were eluted with a 100-ml linear gradient of 50– 500 mM imidazole in Buffer A. Fractions were collected from the peak and their purity was confirmed by 8 % and 12 % sodium dodecyl sulfate-polyacrylamide gel electrophoresis (SDS-PAGE) for RAD52 and RAD52(209), respectively. The pure fractions were loaded onto a Hiload 16/600 superdex column (Cytiva) after equilibration with RAD52 Buffer (50 mM KH_2_PO_4_, pH 8.0, 1 M KCl, 1 mM MgCl_2_ and 0.1 mM EDTA) and RAD52(209) Buffer (10 mM KH_2_PO_4_, pH 7.5, 10 mM NaCl, 1 mM dithiothreitol (DTT), and 0.1 mM EDTA) for RAD52 and RAD52(209), respectively. The fractions were collected from the eluted peak and controlled by an SDS-PAGE for RAD52 and RAD52(209), respectively. Pure fractions were concentrated to approximately 2 mg/ml using an Amicon Ultra 15 centrifugal filter (50K MWCO and 30K MWCO for RAD52 and RAD52(209), respectively) (Merck KgaA, Germany). Protein concentrations were determined using a Nano-Drop 1000 (Thermo Fisher Scientific, Germany). The concentrated proteins were flash frozen in liquid nitrogen and stored at − 80 °C.

### ssDNA binding assays

ssDNA binding was quantified by performing electrophoretic mobility shift assays with ssDNA of different lengths (Table S1). Oligomer sequences were designed for minimal secondary structure formation and purchased from Sigma-Aldrich and Eurofins. Each oligonucleotide was fluorescently labelled at the 3’-end with ATTO565, ATTO680, or ALEXA488. RAD52(209) and RAD52 were diluted to desired concentrations in Buffer 2 (25 mM Tris-Acetate, 2 mM Mg(OAc)_2_, 1 mM DTT, pH 7.5). Proteins were incubated with 10 nM ssDNA either at 25 °C for 10 min, or at 37 °C for 5 min. Afterwards a gel loading dye (New England Biolabs, Cat. # B7024S) was added. Bands were resolved in precooled Tris-acetate-EDTA (TAE) buffer on a 10 % TGX (Tris-Glycine eXtended)-Gel (BioRad). Gels were running at 4 °C and 100 V (≈ 30–35 mA) until the lower front of the gel loading dye approached the bottom of the cassette. Gels were imaged on a Typhoon FLA 9000 (Cytiva). Band intensities were quantified in Fiji^58^. All intensities were normalized to the control value (ssDNA, no protein added). These values were converted to percent and subtracted from 100 %. Final values represent the amount of ssDNA bound to the protein as a function of protein concentration based on the decrease in intensity of bands running at the level of the control ssDNA without protein. Hill equations were fitted to the data weighted with error bars. Fitting constants are given in Table S3.

### Terminated DNA annealing assays

Annealing assays were performed on different pairs of ssDNA (Table S1). Oligonucleotide concentrations were 10 nM. Protein concentrations were 100 nM. RAD52(209) and RAD52 were diluted to the desired concentrations in Buffer 2 and incubated with unlabelled ssDNA at 37 °C for 5 min. Subsequently, complementary 3’-end-fluorescently-labelled ssDNA was added and annealing proceeded at 25 °C. To quantify the annealing rate, reactions were terminated after 1, 3, and 8 min with the STOP-Buffer (5 *μ*M unlabelled ssDNA identical in sequence to the fluorescently labelled one, 10 % SDS, 100 mM EDTA, and 10 units Proteinase K) in a 1:5 STOP-Buffer:reaction-mixture volume ratio and incubated 15 min at 30 °C. Band intensities were quantified in Fiji and normalized to the control value (ssDNA at 0 min). These values were converted to percent and subtracted from 100 %. Final values represent the amount of ssDNA that is annealed at different time points.

To determine the concentration of highest annealing activity, RAD52(209) and RAD52 were diluted in Buffer 2 to desired concentrations in a range from 10 nM to 10 *μ*M. Some protein solutions were preincubated with a 60 nt-long, unlabelled oligonucleotide at 37 °C for 5 min. Afterward a complementary 32 nt-long, 3’-end-ATTO565-labeled oligonucleotide was added. Final oligonucleotide concentrations were 10 nM. The annealing reaction proceeded at 25 °C and was terminated after 15 min as described above. Subsequently, gel loading dye was added. Bands were resolved in precooled TAE buffer on a 10 % TGX-Gel. Gels were running at 4 °C and 100 V (≈ 30–35 mA) for 90 min and imaged using a Typhoon FLA 9000. Band intensities were quantified in Fiji and normalized to the control value (Column C2 in Fig. 5, dsDNA without protein). These values represent the amount of dsDNA product formed at different concentrations.

### Native DNA annealing assay

To visualize intermediate species formed during the annealing reaction, proteins were preincubated at 37 °C for 5 min in Buffer 2 with a 60 nt 3’-ATTO680 labeled oligonucleotide. Subsequently, a complementary, fluorescently labelled oligonucleotide was added (48-ATTO488 or 32-ATTO565). DNA concentrations were 10 nM. Protein concentrations were in the range from 50 nM to 2 *μ*M. Annealing proceeded at 25 °C for 15 min. Reactions were not terminated and immediately loaded on a gel with a non-denaturing gel loading dye (Thermo Fischer, Cat. #R0611). Bands were resolved in precooled TAE buffer on a 10 % TGX-gel. Gels were running at 4 °C and 100 V (≈ 30–35 mA) until the lower front of the gel loading dye approached the bottom of the cassette. As previously described, gels were imaged with a Typhoon FLA 9000 scanner with two excitation wavelengths. Band intensities of both ssDNA and dsDNA were quantified in Fiji. To quantify the amount of unreacted ssDNA, acquired intensities of the ss-DNA were normalized to the corresponding control values of ssDNA (48 nt and 60 nt, respectively). To quantify the amount of non-protein bound dsDNA product, intensities of bands running at the level of the control were normalized to the control value (no protein added). Final values were plotted as functions of protein concentrations.

### Blue native gels

The native oligomeric state of the proteins at micromolar concentrations was visualized in blue native gels. Proteins were diluted in Buffer 2 to the desired concentrations with native detergents (2 % n-dodecyl *β* -D-maltoside (Thermo Fischer, Cat. #BN2005) and 0.5 % digitonin (Thermo Fischer, Cat. #BN2006)). Cathode and anode buffers were prepared according to a standard protocol and kept at 4 °C. Prior to loading, NativePAGE^T*M*^ 0.5 % G-250 Sample Additive (Thermo Fischer, Cat. #BN2004) was added. Bands were resolved on a 4–12 % Novex Tris-Glycine Gel (Invitrogen). Gels were running on ice at 150 V for 1 h and at 250 V for another 2–3 h till the front approached the end of the cassette. NativeMark Unstained Protein Standard (Thermo Fischer, Cat. #LC0725) was used as a reference for molecular weight determination. Gels were fixed, stained with Imperial Blue (Thermo Fischer, Cat. #24615) overnight, and washed thoroughly in water until bands were resolvable. Bands were visualized on an Epson Perfection V700 Photo scanner.

## RESULTS

To determine the oligomeric state of RAD52 as a function of concentration and the influence of its C-terminal domain, we expressed and purified full-length human RAD52, from now on referred to as RAD52, and a version truncated after amino acid 209, referred to as RAD52(209) lacking the RPA and RAD51 binding domains as well as the nuclear localization domain (Fig. 1a, Materials and Methods). With the self-association domain (marked with RAD52 in Fig. 1a) and RAD52’s propensity to form rings, the law of mass action implies that at high concentrations, rings form, which may further interact to aggregates of rings, while at low concentrations, short oligomers are prevalent (Fig. 1b). For simplicity, we define short oligomers here to include monomers, dimers, trimers, and tetramers even though monomers are strictly speaking not oligomers. Using an SDS-PAGE (Fig. 1c), bands for purified RAD52(209) and RAD52 were consistent with their expected monomer molecular weight of 25.3 kDa and 48.4 kDa, respectively.

**Figure 1.**
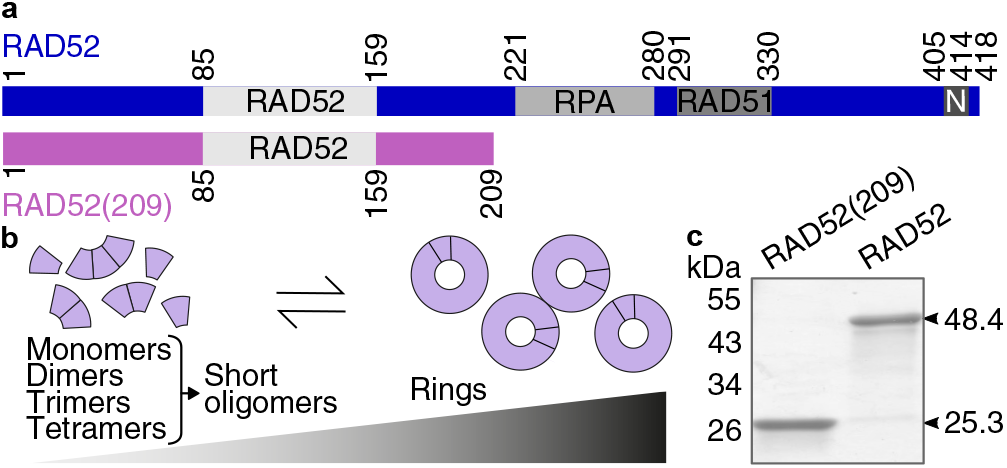
RAD52 domains and multimerization. **(a)** Domain schematic of full-length RAD52 and truncated RAD52(209) including the self-association domain labelled RAD52, the RPA and RAD51 binding domains, and the nuclear localization domain N. **(b)** The oligomeric state depends on the protein concentration. Higher concentrations shift the equilibrium oligomeric state from short oligomers to rings and their aggregates. **(c)** SDS-PAGE of purified RAD52 and RAD52(209) with indicated sequence mass.

### Short RAD52 oligomers and not rings dominate at physiological concentrations

To measure the oligomeric state of individual complexes under native conditions, we used mass photometry^57^—a label-free, single-molecule technique that measures the molecular weight of individual proteins or complexes based on the interference contrast from light scattered back from surface-adsorbed molecules. The molecular-weight measurement and inferred RAD52 stoichiometry show that the abundance of short oligomers decreases and that of rings and their aggregates increases with increasing protein concentrations (Fig. 2). Both protein mass histograms have peaks corresponding to short oligomers and rings with the relative abundance changing with concentration as expected from the law of mass action (Fig. 2a,b; Figs. S1 and S2 for recordings on various days). With increasing concentration, there was also a small number of large aggregates with megadalton molecular weights (not shown in Fig. 2; for an example, see inset top row Fig. 7). Also with increasing concentration, monomers multimerize to dimers, trimers, or tetramers (Fig. 3a, Table S2). Please note though that we could only detect monomers of the full-length protein because the molecular weight of RAD52(209) was below the 40 kDa-lower-mass-detection limit of the mass photometry instrument we used.

**Figure 2.**
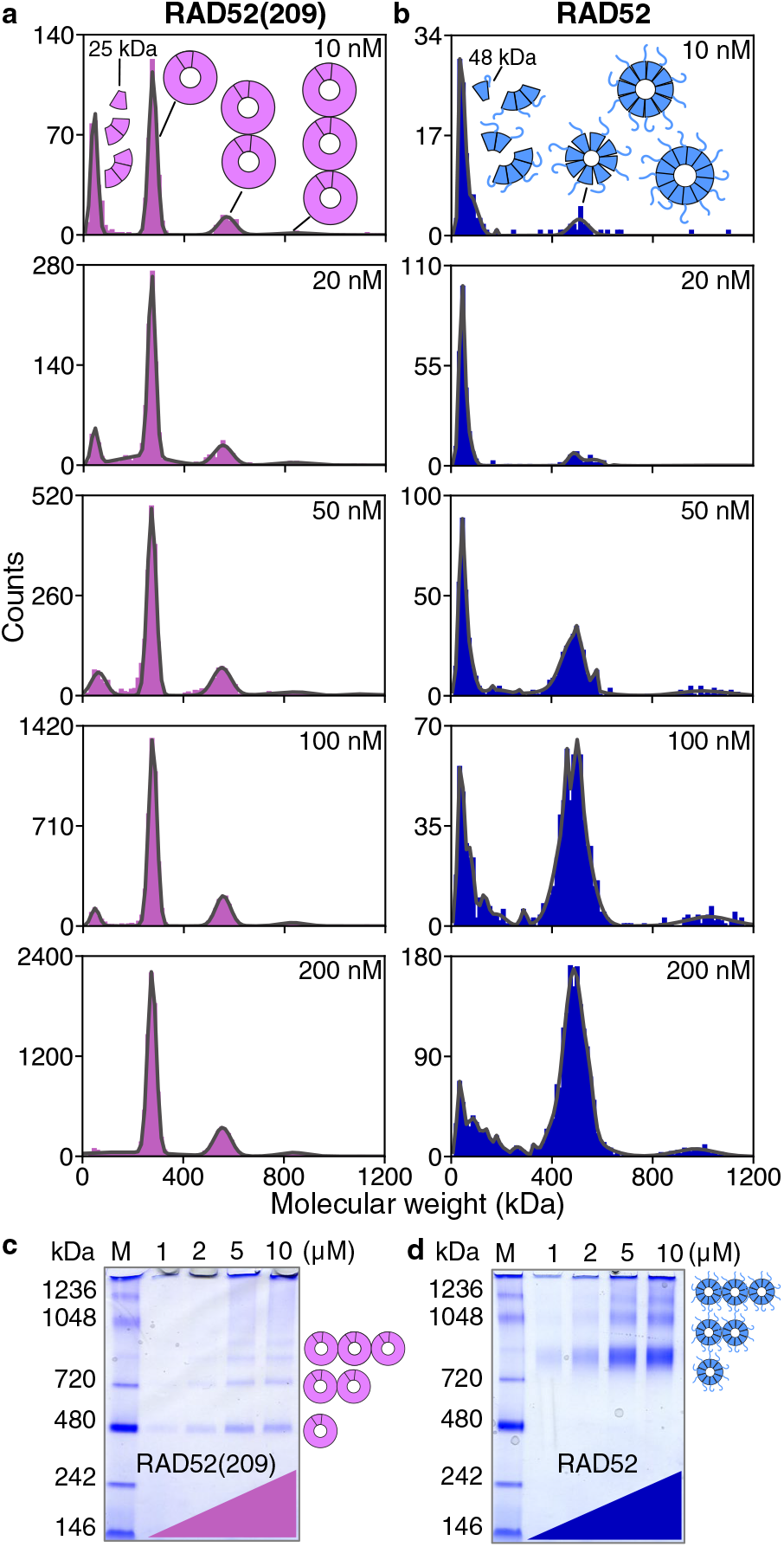
Mass-photometry molecular-weight histograms of **(a)** RAD52(209) and **(b)** RAD52 for increasing concentrations indicated in the panels. Lines are fits of multiple Gaussians. Each dataset is a cumulative distribution from four measurements on different days. Schematics next to or above peaks illustrate the peak’s composition of short oligomers or rings. Blue native gels of **(c)** RAD52(209) and **(d)** RAD52 at higher concentrations show aggregates of rings (M: marker).

**Figure 3.**
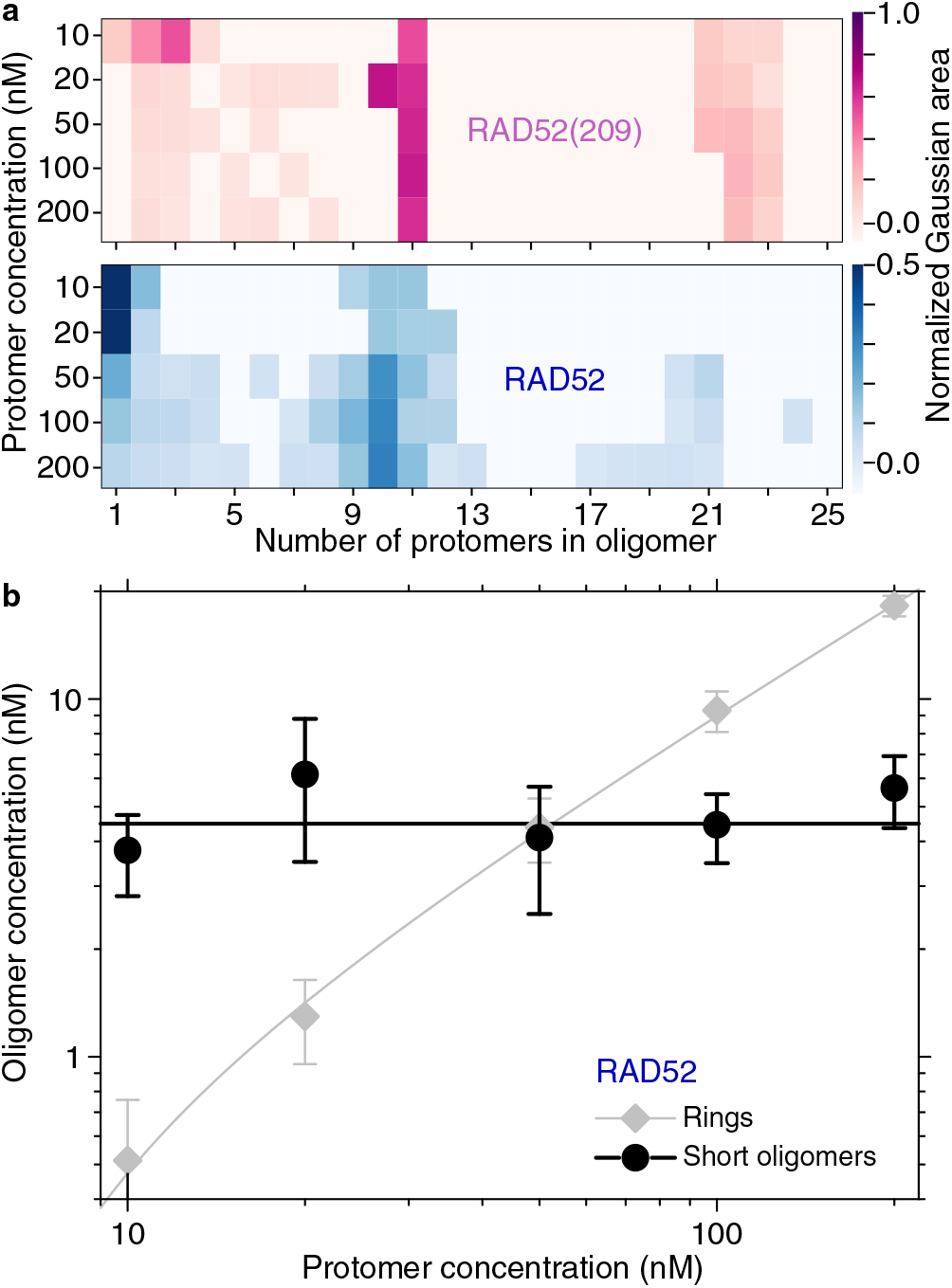
Abundance of RAD52 oligmers. **(a)** Normalized abundance of oligomers as a function of concentration. The color scales on the right represent the normalized area under each oligomer in Fig. 2. **(b)** Concentration of RAD52 short oligomers and rings (sums of 1–4 mer and 8–12mer concentrations, respectively) based on their counts in Fig. 2b as a function of the protomer concentration. Lines are fits to the data (see text).

Peaks associated with rings are prominent and there is a gap in oligomer abundance between short oligomers and rings (Fig. 2a,b, Fig. 3). This gap implies that rings are thermodynamically very stable and that ring closure is associated with an extra energetic benefit^59^. For RAD52(209), more rings were present at lower concentrations implying that either self-association or ring closure is stronger compared to RAD52. For RAD52, we detected a higher relative abundance of short oligomers compared to RAD52(209). For RAD52 at the lowest measured concentration of 10 nM, only few rings were present (Fig. 2b, Fig. 3a). When we determine the concentrations of the various RAD52 oligomers based on the counts, the analysis showed that there was nearly a 10-fold excess of short oligomers over rings (Fig. 3b). Short oligomers were mostly monomers and some dimers with trimers and tetramers forming above 20 nM (Fig. 2, Fig. 3a, and Figs. S2). As a function of protomer concentration *c*_*protomer*_, the short oligomer concentration was on average constant (4.5 *±* 0.4 nM, mean *±* fit error, black line in Fig. 3b) and the ring concentration *c*_*ring*_ increased in a linear fashion according to *c*_*ring*_ = (*c*_*protomer*_ − *c*_*crit*_)*/n*_*avg*_ with the critical concentration *c*_*crit*_ for ring formation of 5.0 *±* 0.7 nM and the average number of protomers per ring *n*_*avg*_ of 10.6 *±* 0.2 (gray line in Fig. 3b). Thus, according to the fit, a minimum RAD52 concentration of about 5 nM is required for ring formation. Summarizing, the most abundant RAD52 oligomeric species at estimated physiological concentrations are likely to be monomers and dimers (Fig. 3b).

Rings had a variable quaternary structure and abundance between the two RAD52 variants (Fig. 2a,b and Fig. 3a). As expected, RAD52(209) formed undecameric rings with a molecular weight of 279.6 *±* 1.3 kDa (mean *±* SEM based on fit error of the mean) and aggregates of rings (Fig. 2a). This molecular weight is in excellent agreement with the calculated one of 278.3 kDa. The undecamer peak’s standard deviation was about 18 kDa, smaller than those of proteins used for calibration (for example, peaks at 146 *±* 36 kDa and 480 *±* 47 kDa [mean *±* SD based on Gaussian fit]^57^). The small undecamer-peak standard deviation indicates that the peak width is due to the measurement error and does not arise from a variation in stoichiometry. Surprisingly, RAD52 did not solely form heptamers^19,20^ or undecamers^21,22^. The ring stoichiometry varied continuously between 7–13 protomers, with decamers having the highest abundance consistent with the value of *n*_*avg*_ determined above (Fig. 2b and Fig. 3a). Interestingly, RAD52 did not show significant ring aggregation [i.e. peaks at two and three times the ring molecular weight were smaller compared to RAD52(209)] at nanomolar concentrations contrary to the finding that the C-terminal domain should promote aggregation^19^.

At micromolar concentrations, rings of both RAD52 variants formed aggregates (Fig. 2c,d). Blue native gels showed prominent bands that we attribute to rings and their aggregates with the lowest-molecular weight band corresponding to a single ring. Bands associated with aggregates of rings decreased in intensity with larger aggregate size. Both proteins ran at higher molecular weights than predicted as observed previously^60^. Furthermore, RAD52 ring bands were smeared out and more intense compared to the RAD52(209) ones. These observations are consistent with an inhomogenous ring quaternary structure of RAD52 and its C-terminal domain promoting aggregation at micromolar concentrations^19^, respectively. Thus, based on the differences between RAD52 and RAD52(209) in both assays, the C-terminal domain appears to affect ring aggregation, ring closure and stoichiometry, and the self-association of RAD52.

### RAD52 binds cooperatively to ssDNA

To verify the functionality of the proteins and measure their binding affinities to DNA, we used electrophoretic mobility shift assays (Fig. 4). First, we tested the ability of both proteins to bind to 32 nucleotide (nt)-long ssDNA at 37 °C. At low protein concentration, there were only small shifts of the ssDNA bands indicating that there were either no or weak, transient interactions with the proteins (insets Fig. 4a). With increasing protein concentration, ssDNA bound to the proteins and migrated slower. However, shifts did not result in discrete bands but in a smear on the gels indicating transient interactions of various strengths and/or different sizes of associated proteins. We determined the amount of protein-bound ssDNA as a function of concentration (Materials and Methods) and fitted a Hill equation (solid lines in Fig. 4a) to the data. The *K*_*d*_-values were about 250 nM and 135 nM for RAD52(209) and RAD52, respectively, indicating about a two-fold higher ssDNA binding affinity for RAD52 (see Table S3 for fit parameters). The Hill coefficients were 3.1 *±* 0.5 and 2.7*±* 0.2 for RAD52(209) and RAD52, respectively. These coefficients are significantly different from 1 (p-value of 0.004 and less than 0.0001, respectively) implying cooperative binding of a few monomers^61^. Instead of short-oligomer binding, ring binding would have resulted in a steep rise in the bound fraction with a very high cooperativity, i.e. with the Hill coefficient on the order of 10, which we did not observe, indicating that the rings are not the preferential substrate for protein-DNA interaction^61^. As a reference, the dashed and dotted lines in Fig. 4a show how non-cooperative and ring binding curves with the same *K*_*d*_ as the one for RAD52 look like (Hill coefficient of one [Michaelis-Menten] and 11 [rings], respectively). Thus, our ssDNA binding curves are *inconsistent* with a ssDNA-ring-binding mechanism, but more *consistent* with short oligomers binding sequentially or independently^61,62^. Sequential binding—i.e. monomers and dimers bind one after another to form a trimer rather than a trimer binding directly— may explain the systematic deviation of the data from the Hill equation at low RAD52 concentrations in Fig. 4a. While Hill-equation fit parameters were similar for longer ssDNA, experiments performed at room temperature showed a lower cooperativity (Fig. S3 and Table S3). To summarize, both RAD52(209) and RAD52 bind ssDNA cooperatively. Co-operativity means that once a monomer or short oligomer is bound, the affinity for further binding is increased. Since allostery is unlikely to be present for non-neighboring binding sites on the oligonucleotide, the cooperativity implies that RAD52 molecules bind next to each other forming a short nucleoprotein filament.

**Figure 4.**
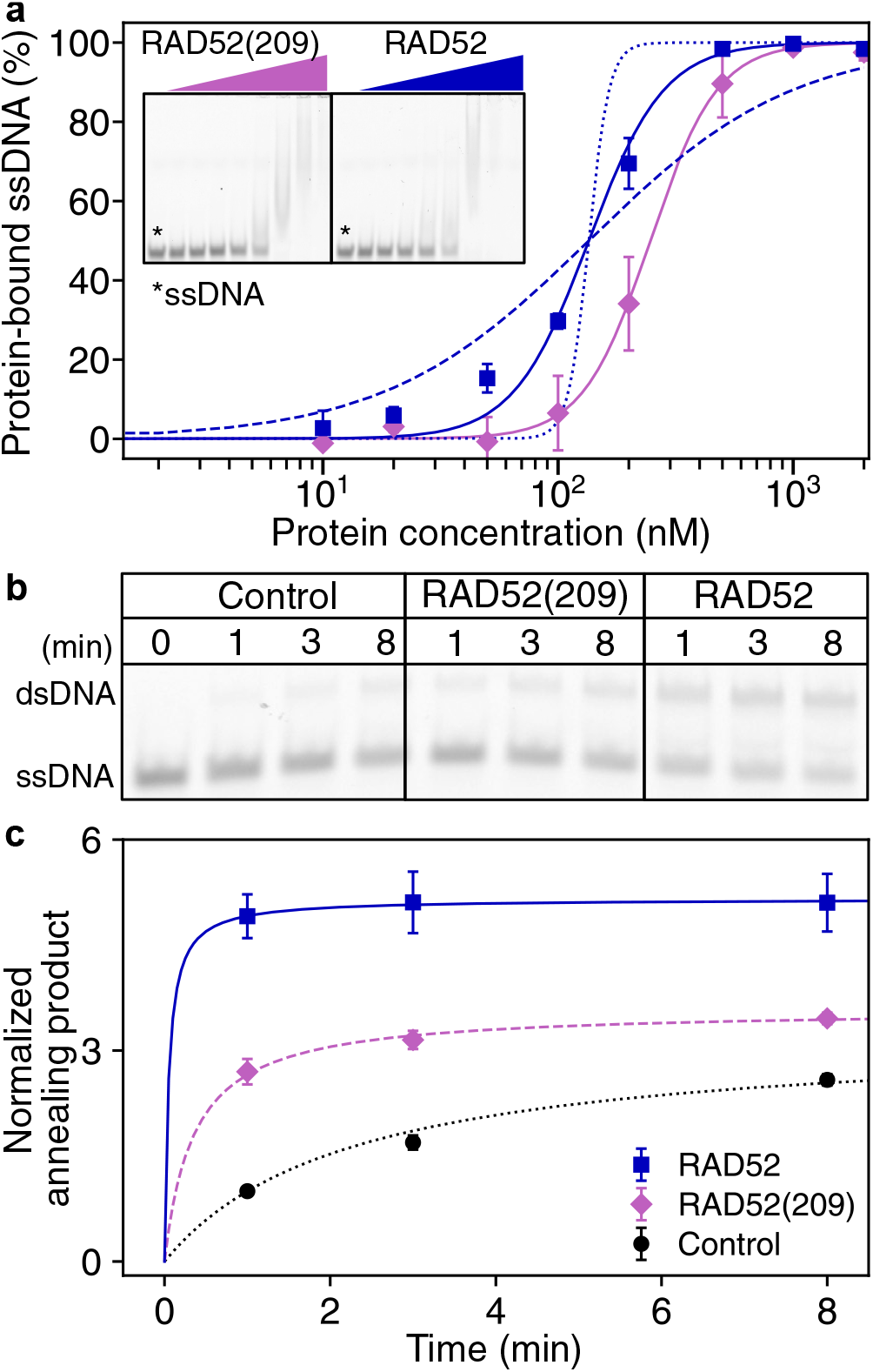
RAD52 single-strand DNA binding and annealing assays. **(a)** Amount of protein-bound ssDNA (10 nM, 32 nt long at 37 °C, mean *±* SEM from 3 gels) as a function of concentration. Solid lines are Hill equation fits (see Table S3 for fit parameter values). Dashed and dotted lines are plots of the Hill equation with a fixed *K*_*d*_ and Hill coefficient of 1 and 11, respectively. An exemplary binding gel is shown in the inset with the first column being a control (no protein, marked with a star) and the others correspond to the plotted concentrations. **(b)** Exemplary annealing gel using complementary 32 nt labeled 3’-ATTO680 and unlabeled 60 nt oligonucleotides for the annealing reaction as a function of time (Control: no protein). Protein and DNA concentrations were 100 nM and 10 nM, respectively. **(c)** Annealing product as a function of time (mean *±* SEM, 3 repeats). Lines are fits for second-order reaction kinetics.

**Figure 5.**
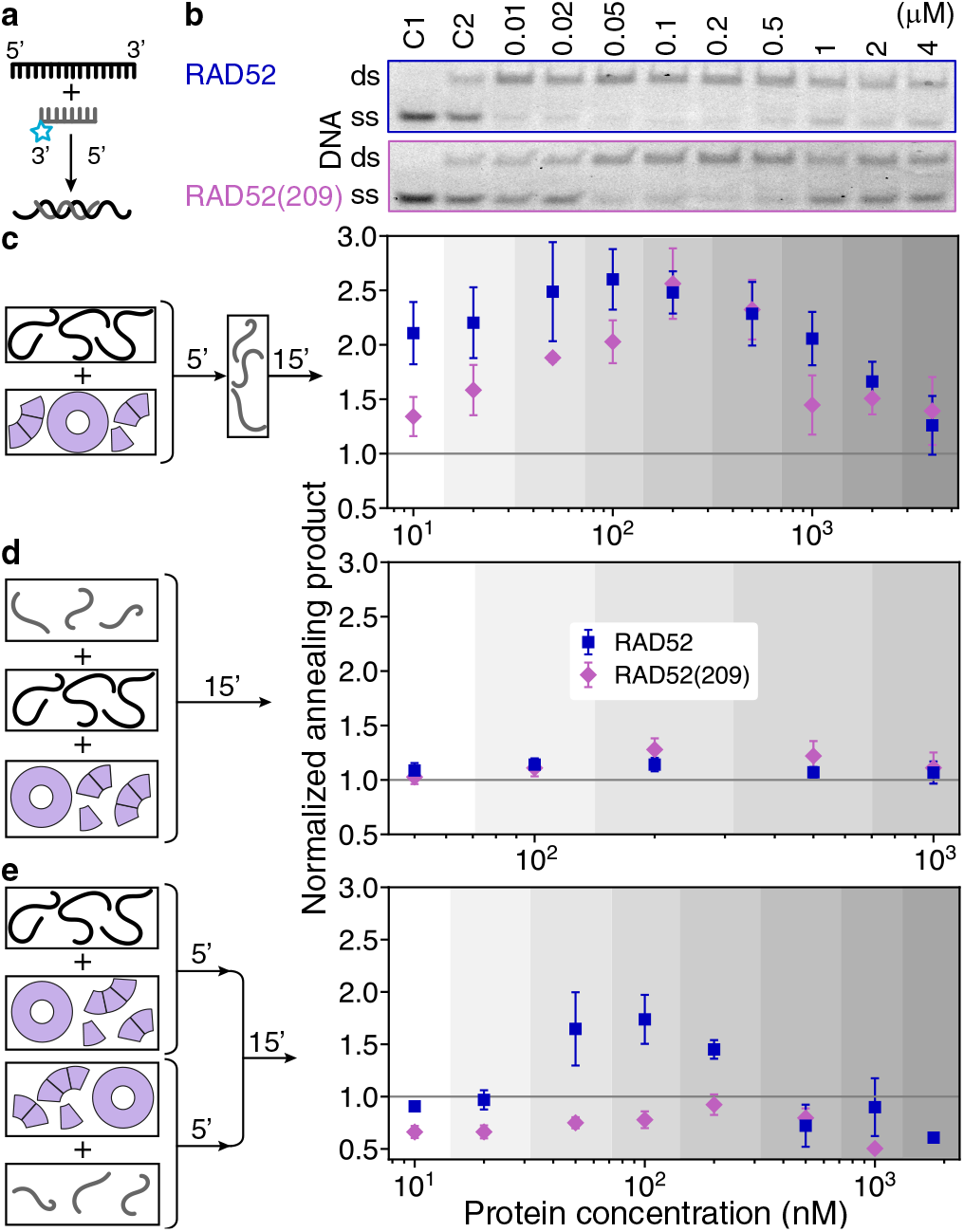
dsDNA product yield of the annealing reaction. **(a)** Hybridizing-reaction schematic of a 32-nt long, fluorescently-labeled (gray with cyan star) and a 60-nt long, unlabeled (black) oligonucleotide without protein. **(b)** Annealing gels of electrophoretic mobility shift assays when one strand is incubated with the protein before mixing it with the other strand. Control C1: only one, the labeled oligonucleotide. C2: annealing control, i.e. hybridizing reaction of both oligonucleotides without protein. Schematic and quantification of **(c)** the annealing reaction from (b) with *one strand incubated with the protein before adding the other strand*, **(d)** the annealing reaction *with both strands and the protein mixed at the same time*, and **(e)** the annealing reaction with *both strands incubated separately with the protein before mixing*. Incubation times in min are indicated on the arrows. All data points are mean values *±* SEM (6 repeats).

### High RAD52 concentrations promote ring binding and inhibit annealing

To determine annealing rates and efficiencies, we incubated complementary strands with the single-strand annealing proteins and varied the protein concentration and the order of mixing and incubation of proteins with individual strands (Fig. 4b,c and Fig. 5, see Materials and Methods). With a protein excess (100 nM) over DNA (10 nM) and incubation of only one strand with the proteins before adding the complementary strand, both RAD52(209) and RAD52 promoted annealing. With increasing annealing time, dsDNA bands of electrophoretic mobility shift assays became more intense (Fig. 4b). After quantifying band intensities, we could fit a second-order rate equation to the annealing data consistent with previous observations on the reaction kinetics^1^ (Fig. 4c). Both RAD52(209) and RAD52 increased the annealing rate compared to the control of hybridization without protein about 6- and 30-fold, respectively (Table S3).

Next, we determined how the dsDNA product yield of the annealing reaction depended on concentration and incubation order (Fig. 5). We normalized all annealing product band intensities by the corresponding hybridizing reaction without protein (Fig. 5a and C2 in Fig. 5b). When we incubated only one of the two strands with the protein before adding the other strand, annealing product yield was promoted up to a 2.5-times at about 100–200 nM for both RAD52 variants (Fig. 5b,c).

RAD52 was more efficient in annealing at lower concentrations compared to RAD52(209). In particular, the annealing product formation was already doubled at 10 nM compared to the hybridizing reaction without protein. At this concentration, hardly any rings and mostly shorter oligomers were present for RAD52 (Fig. 3b). By contrast, with increasing concentrations above 1 *μ*M, the amount of annealed dsDNA product and the amount of the unreacted ssDNA (Fig. 5b) almost reached control levels without protein. Thus, under these conditions ssDNA was bound to non-annealing-competent RAD52 complexes.

When the two complementary strands were not incubated with the protein, but both directly mixed with the protein, annealing was not promoted much at any protein concentration (Fig. 5d). Thus, there was not more product within the 15 min reaction time compared to the hybridizing reaction without protein. This finding suggests that initial nucleoprotein filament formation for homology search is as diffusion limited as hybridization.

When both strands were incubated with protein before mixing, annealing was not promoted much or even inhibited (Fig. 5e). Inhibition was particularly strong for RAD52(209) for which product formation was cut in half at 1 *μ*M. Only for RAD52, annealing was promoted at 50–200 nM but not as much compared to the reaction when one strand was incubated with protein before adding the other strand. Since annealing was not promoted for RAD52(209) at these concentrations, RAD52’s ability to promote annealing under these conditions must be due to the C-terminal domain. Inhibition is consistent with earlier observations^51^ and suggests that for recombination and homology search only one strand should have an annealing-competent RAD52 oligomer bound and the complementary strand not.

To gain insight into the annealing-competent and inhibiting oligomeric species, we used fluorescent labels on both strands and our most efficient annealing conditions (Fig. 6a). Also, we did not stop the annealing reaction. Individual bands of the oligonucleotides and the dsDNA product are visible at the bottom of the gel in Fig. 6a. For 50–200 nM RAD52(209), the dsDNA ran at the same level as the control. Thus, there was no protein bound. However, there was more product compared to the negative control (solid diamonds in Fig. 6b with values larger than one). This finding suggests that the protein-mediated ssDNA annealing leads to the unbinding of the protein and that interactions of the DNA with the proteins were transient. With increasing concentration, higher bands appeared that we attribute to DNA-bound rings. Since gels ran for about one hour, these DNA-bound rings were stable complexes. At these concentrations, we also observed the decline in unbound product yield (compare with Fig. 5).

**Figure 6.**
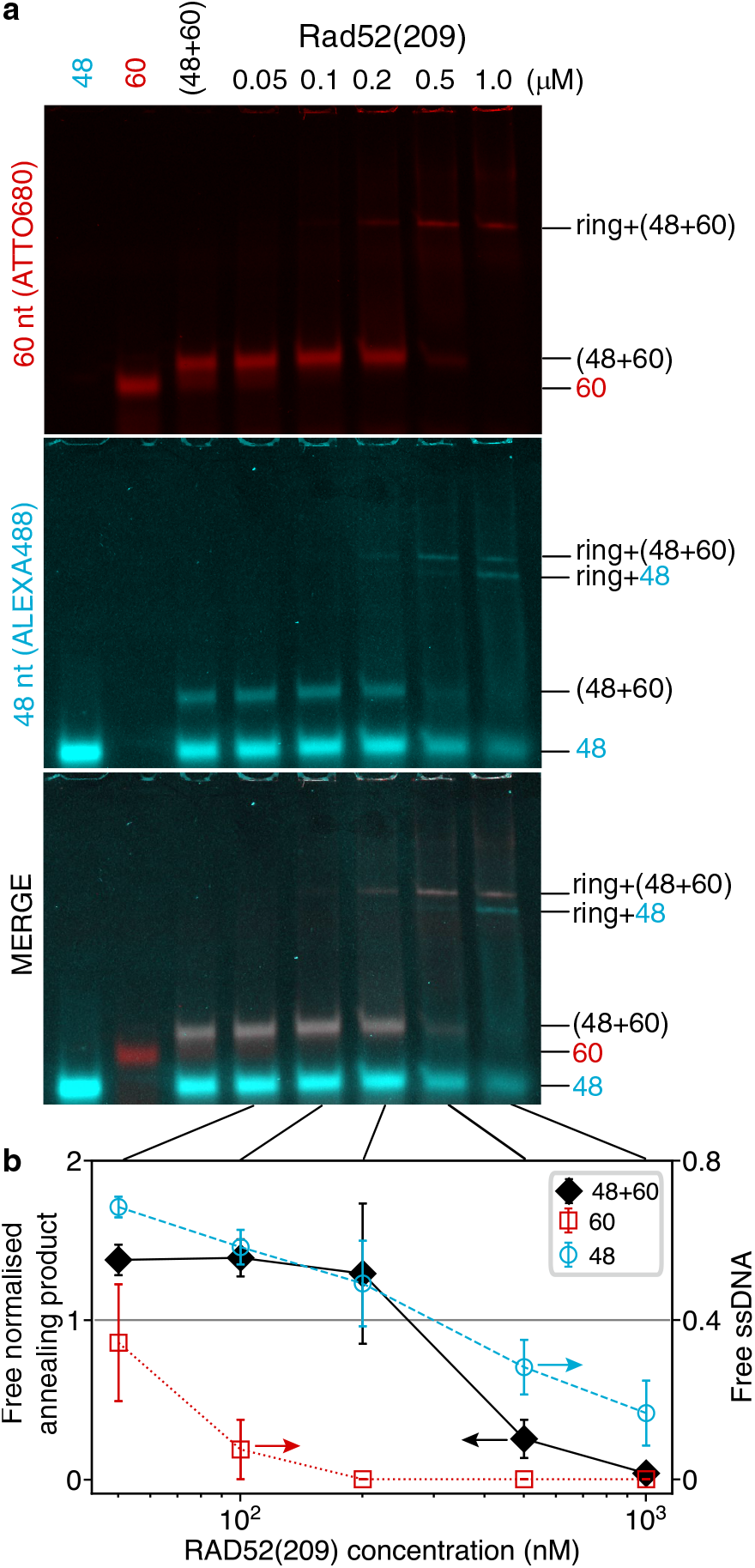
Annealing gels and product yield of fluorescently-labeled oligonucleotides. **(a)** Native gels of complementary oligonucleotides with increasing concentrations of RAD52(209) (the 60 nt-long strand incubated with the protein before adding the other strand). The three controls on the left are without protein. **(b)** Amount of annealing product (solid black diamonds) and unreacted ssDNA (open cyan circles and red squares) normalized by control values plotted as a function of RAD52(209) concentration (mean *±* SEM, 6 repeats).

At the highest tested protein concentration of 1 *μ*M there were two separate “ring” bands—one containing both oligonucleotides and the other lower band containing only one strand (the shorter one, cyan in Fig. 6a). That DNA-bound rings ran at different levels implies that they had different mobilities. Such differences may arise from a different length and charges of the oligonucleotides. Whether the two strands, in the band that contains both oligonucleotides, were also annealed or bound separately on different or the same ring is unclear. Summarizing, at 1 *μ*M nearly all DNA was bound to rings or larger aggregates. Since there was already a significant decline in the annealing activity at this concentration, these findings suggests that neither ring-nor aggregate-bound DNA are beneficial for annealing.

### ssDNA binds and aggregates RAD52 rings

To determine on the single-molecule level, to which oligomeric RAD52(209) and RAD52 species ssDNA binds, we monitored the annealing reaction via mass photometry under our most efficient annealing conditions (Fig. 7). After incubation of the first oligonucleotide (10 nM, 60 nt, 18.6 kDa, no fluorescent label) with the proteins (100 nM), the number of rings decreased and their molecular weight was shifted (middle row Fig. 7). The abundance and molecular weight of short oligomers were unaffected. The well-defined, RAD52(209) single- and double-ring peaks shifted by 42 *±* 1 kDa and 74 *±* 4 kDa, respectively, corresponding to about two and four bound ssDNA with a combined theoretical molecular weight of 37 kDa and 74 kDa, respectively (Fig. 7a and Fig. S4). As with the annealing gel assay (Fig. 6), it is unclear to which binding site on the ring DNA was bound. Furthermore, it is important to stress that two identical ssDNAs were bound to the same ring. Such complexes are hard to reconcile with proposed ring-annealing models^2,16,27,33–35^. With shorter oligonucleotides, the shifts were also consistent with two bound strands (Fig. S4). For longer oligonucleotides that can wrap around the ring twice, only about 1.5 strands were bound on average to single rings (Fig. S4). For RAD52, molecular-weight shifts were difficult to quantify due to the spectrum of ring structures (Fig. 7b).

We attribute the reduction in ring counts to DNA-protein aggregates. First, we observed an increased count of higher molecular-weight species (for example, a peak at ≈ 3 MDa for RAD52(209), insets Fig. 7). In addition to these species, larger aggregates were visible in the raw images but were beyond the upper mass-calibration limit of around 6–7 MDa (Fig. S5). Since lower oligomer counts did not change much, aggregates must have been largely composed of rings. We emphasize that mass photometry is representative of object concentration, meaning that these large aggregates occupied a significant fraction of total RAD52 present in solution.

Addition of the complementary strand (10 nM, 32 nt, 9.8 kDa, no fluorescent label) reduced ring counts further but did not change the molecular-weight shift of ring peaks (bottom row Fig. 7, Table S4). Ring aggregates were still present. RAD52(209) single-ring peaks shifted by 42 *±* 2 kDa not significantly different from the shift when incubated with the first single strand. The dsDNA product’s molecular weight was below the detection limit. A lower molecular weight nucleoprotein filament with annealed DNA was also not detected. Thus, while ring counts were affected, mass photometry could not detect a clear signature of the annealing reaction. Taken together, we can conclude for the ssDNA-protein interaction that (i) two identical ssDNAs can bind a single ring, (ii) ss-DNA can lead to RAD52(209) and RAD52 aggregate formation, and (iii) the detected fraction of short oligomers did not bind ssDNA and change abundance on normal, cleaned glass surfaces.

Since glass surfaces are negatively charged, we wondered whether the negatively charged DNA bound to RAD52 may decrease the binding affinity of such a complex to the surface. To test this hypothesis, we used a modified, positively charged glass surface for mass photometry^63^. On positively charged surfaces using protein incubated with ssDNA in the same manner as in Fig. 7, complexes bound rigidly and we could indeed detect a shift in molecular weight not only for rings but also for short oligomers (Fig. 8). The first peak in the mass histogram of Fig. 8a is consistent with the molecular weight of a RAD52 monomer without ssDNA. The second peak, we attribute to a monomer bound to ssDNA. This peak’s molecular weight is significantly different from the sequence molecular weight of a dimer without DNA (dashed vertical line marked with a two). For subsequent peaks, we assigned shifts in consecutive order to RAD52 oligomers. In this manner, the shifted decamer peak still had the highest abundance comparable to the data without ssDNA (Fig. 3a). Shifts were consistent with two single strands bound per short oligomer up to heptamers and an increasing number of strands—up to seven ssDNA—bound to single rings (Fig. 8b). This number is approaching estimates of previous experiments, in which up to 12 ssDNA strands could have been bound to a horseshoe-shaped complex—not a ring^21^. More than two bound strands suggests that some complexes have DNA strands that do not fully wrap around rings, but have parts dangling freely from the rings. Such DNA tethers may increase the negative charge of the complexes and prevent their binding to normal, negatively charged glass surfaces for which we detected only rings with two bound ssDNA for RAD52(209). Together this data shows that short oligomers including monomers can bind two strands.

**Figure 7.**
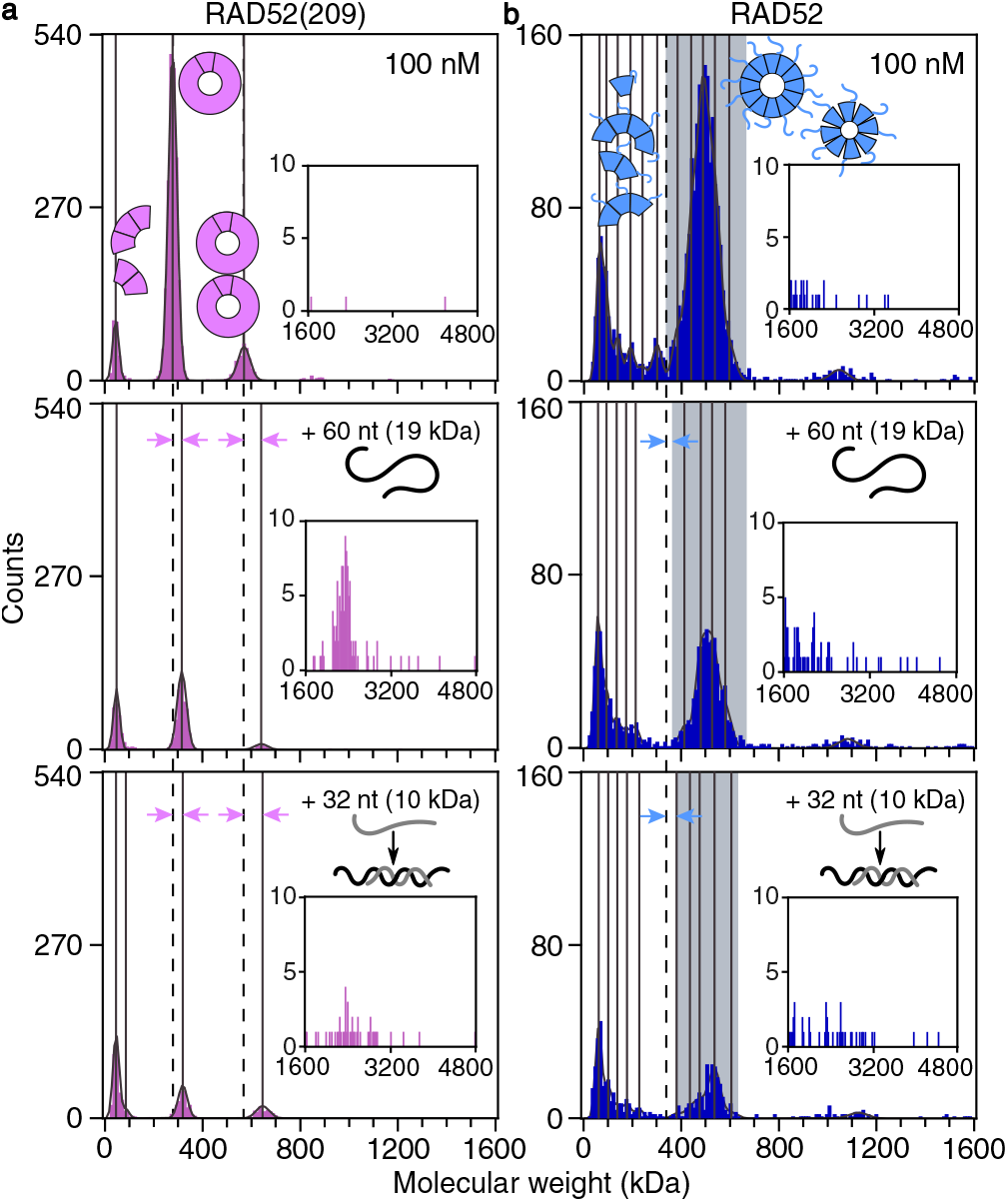
Annealing observed with mass photometry. **(a)** RAD52(209) and **(b)** RAD52 without DNA (both proteins 100 nM, top row), in presence of ssDNA (10 nM, 60 nt, unlabelled, middle row), and after the addition of the complementary strand (10 nM, 32 nt, unlabelled, bottom row). Vertical solid lines indicate peak centers. Dashed lines mark RAD52(209) peak centers or the start of the RAD52 ring peak without added DNA. Molecular weight shifts upon DNA binding correspond to distance between arrowheads. Each dataset is a cumulative distribution from four measurements.

**Figure 8.**
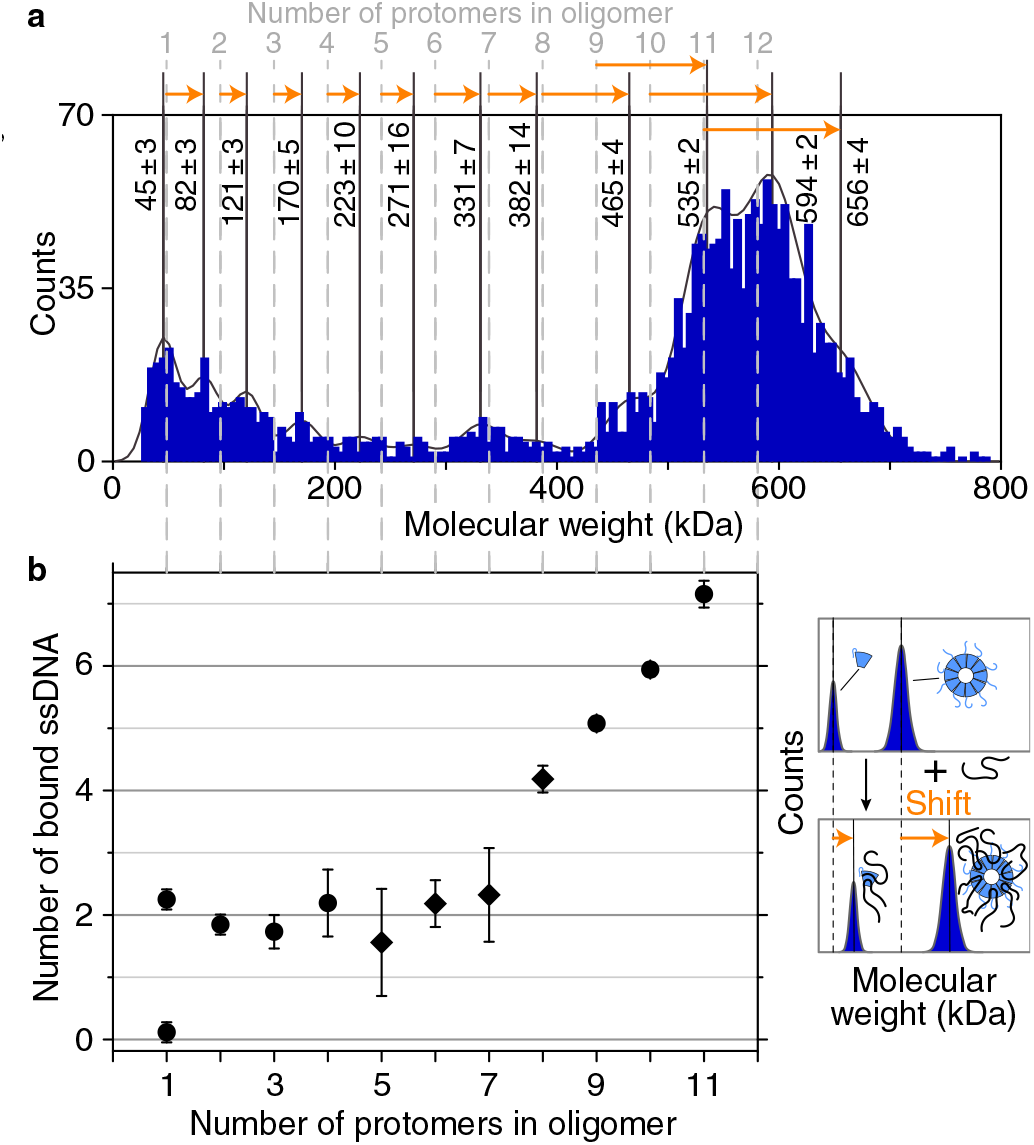
DNA binding with mass photometry on positively charged surfaces. **(a)** Mass histogram of RAD52 incubated with ssDNA on an APTES-coated surface^63^ (100 nM protein and 100 nM unlabeled 60 nt-long, 18.6 kDa ssDNA). Dashed vertical lines correspond to the sequence-based mass of RAD52 oligomers and solid lines to detected peaks with marked molecular weights (centers *±* SEM). **(b)** Molecular weight shift of peaks (orange arrows in **(a)** illustrated to the right) relative to measurements (circles) or the sequence-based mass (diamonds) in the absence of ssDNA (Table S2) normalized to the molecular weight and different mass sensitivity^63^ of ssDNA (a 7 % correction). Error bars correspond to SEMs of **(a)**.

### ssDNA excess disassembles rings and aggregates

To test whether more ssDNA could disassemble rings as previously suggested^21^, we increased the ratio of ssDNA to protein using standard, negatively charged glass surfaces (Fig. 9). With 100 nM of protein and adding increasing amounts of ssDNA (10 nM, 100 nM or 200 nM), the molecular-weight shift of the detected ring peak did not significantly change with more ssDNA present. There were on average about two oligonucleotides bound as with the previous annealing experiments on negatively charged surfaces (Fig. 7). Again we did not observe a shift for short oligomers but a more than 10-fold increase in counts of short oligomers at the highest used ssDNA concentration (bottom row in Fig. 9). Also, the counts of larger species and aggregates decreased; and, for RAD52(209), the number of rings increased. We attribute the increase in ring count to a disassembly of larger species and aggregates. The increase in the number of short oligomers suggests that rings were disassembled with an excess of ss-DNA consistent with previous work^21^ and in the favor of a mechanism that uses short oligomers to anneal ssDNA.

**Figure 9.**
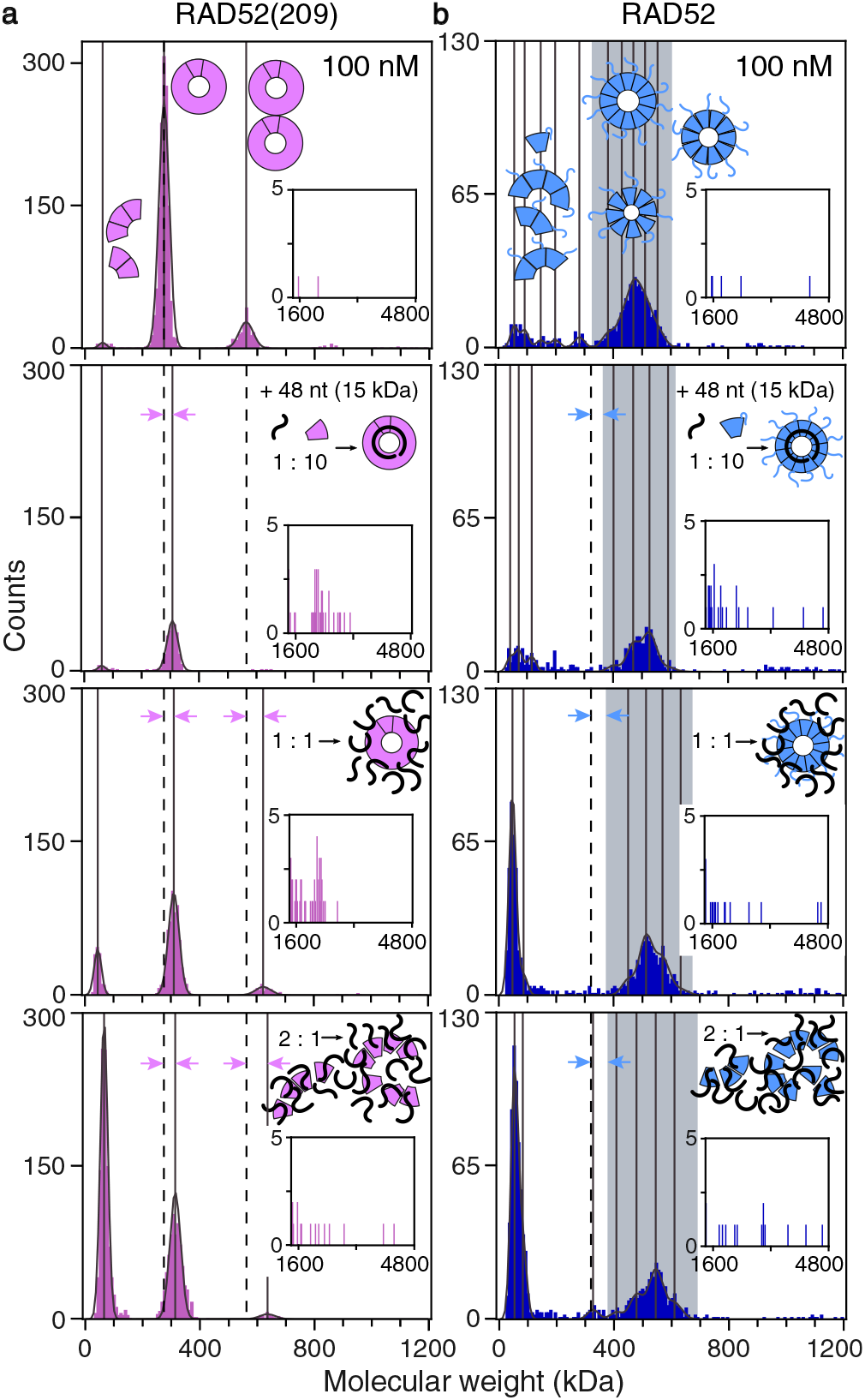
Mass photometry with an excess of DNA over protein. Mass histograms of **(a)** RAD52(209) and **(b)** RAD52 (both 100 nM and on normal glass surfaces) before (top row) and after the addition of ssDNA (unlabeled 48 nt; 10 nM, 100 nM, and 200 nM, respective following rows). Different protein-to-DNA ratios are illustrated as insets. Vertical solid lines indicate peak centers. Dashed lines mark RAD52(209) peak centers or the start of the RAD52 ring peak without added DNA. Molecular weight shifts upon DNA binding correspond to distance between arrowheads. Each dataset is a cumulative distribution of three measurements.

## DISCUSSION

### RAD52 monomers dominate over rings at physiological concentrations

Mass photometry enables the measurement of molecular weight and oligomerization state of native complexes at nanomolar concentrations corresponding to the estimated *in vivo* RAD52 concentration. Our data showed that at these physiological concentrations, even up to about 50 nM, the RAD52 population predominantly consisted of monomers and short oligomers only (Fig. 2b and Fig. 3). Oligomerization and ring formation may be understood in terms of isodesmic growth^64^ in combination with a cyclization model^59^. Based on such a model, rings form and their concentration increases in a near linear fashion above a certain critical concentration while the monomer concentration remains nearly constant^59^. This concentration dependence of short oligomers and rings is consistent with our data (Fig. 3) and similar to that of the phage RAD52 homologue, Red*β* ^37^. Our measured critical concentration of 5 nM for RAD52 ring formation is larger than the estimated *in vivo* concentration of 2 nM in humans^55^. Thus, based on our data and theory, RAD52 short oligomers dominate over rings—if any are present at all—at estimated physiological concentrations. Based on abundance, this data suggests that the annealing-active form of RAD52 by itself may be monomers or short oligomers similar to its active form in ssDNA-RPA-Rad52 complexes^21^ or its homologue Red*β* ^37^.

### RAD52 forms rings with a variable number of protomers

Consistent with the literature^15^, we found that C-terminally truncated RAD52(209) mostly formed undecameric rings (Fig. 2a). For full length RAD52, by contrast, we observed a spectrum of rings with a variable number of protomers from 7–13, with the decamer in highest abundance (Fig. 2b and Fig. 3a). When RAD52 was described to be a heptamer, the authors also noted the ring heterogeneity and broad width of its mass distribution^20^, which is consistent with a variety of rings. Other early measurements also indicated that RAD52 complexes may contain 4–13 protomers^19^ or were “amorphous in shape” and variable in size with only occasional rings being visible^65^. However, hydroxyl radical treatment of ssDNA-RAD52 complexes revealed 4-nt binding per monomer and repeats up to 36–40 nt consistent with a maximum number of 9–10 protomers per ring^28^. Also, recent negative stain electron microscopy imaging could not resolve any distinct seven-fold rings (Fig. S3 in Stefanovie *et al*.^66^). The latest cryo-EM data shows that RAD52 can form undecameric rings. Whether there were other stoichiometries present is unclear as the undecameric structure from C-terminally truncated RAD52 was used as a template for classification^22^. Our data also showed less frequent and heterogeneous RAD52 ring formation suggesting that the presence of the C-terminal region could play an auto-inhibitory role in ring formation. Taken together, our findings are consistent with previous reports and show that RAD52 ring stoichiometry is variable. Thus, interpretations of data in terms of heptameric or other static ring models that rely on a fixed stoichiometry have to be revisited^35^.

### RAD52 binds ssDNA cooperatively

We found that RAD52 binds ssDNA cooperatively with a Hill coefficient of about three (Fig. 4a). A moderate cooperativity suggests that short oligomers assemble to a short nucleoprotein filament containing at least a RAD52 dimer. That short oligomers can bind ssDNA is directly supported by our mass photometry data on positively charged surfaces (Fig. 8). Presumably, nucleoprotein filaments should be nucleated at a 3’ end because of RAD52’s end binding preference protecting ends against exonucleases^28,65^. A short filament consisting of a few RAD52 molecules may be the functional entity to effectively search for homology^38^. Interestingly, cooperativity increased with increasing temperature (Fig. S3 and Table S3) suggesting that ssDNA binding and nucleoprotein filament formation requires the transition over an activation barrier.

With an excess of ssDNA over protein (Fig. 9), we observed less DNA-RAD52 aggregates and more short oligomers, the latter finding being consistent with earlier observations^21^. The increase in short-oligomer abundance suggests a dissociation of rings. Ring dissociation may be explained by the thermodynamics of DNA-RAD52 binding. Initial binding is endothermic^27^ and only driven by a large gain in entropy. The gain in entropy may be due to the dissociation of the rings resulting in many short oligomers being able to bind to many sites on the ssDNA^67^. In essence, there are many more states that individual monomers of a dissociated ring can have with a ssDNA strand compared to the single state of a strand being bound to a ring. Thus, entropy may favor dissociation of the rings and could suggest that rings serve as a protein reservoir. Yet, in principle, both rings and small oligomers bind ssDNA and could be the substrate for subsequent homology search and annealing.

### Competing reactions during annealing

To understand enhancement and inhibition, it is helpful to look at the two extreme conditions of low and high protein concentration. In these limits, either RAD52 monomers or rings dominate in terms of their abundance. While inhibition at low protein concentrations may reflect function in terms of insufficient complementarity for homology search, inhibition at high concentration may be a consequence of RAD52’s biochemistry. Competing, concentration-dependent, biochemical reactions during annealing include: (i) RAD52 oligomerization, (ii) ring closure, (iii) ring self-association and/or large aggregate formation/phase separation^68^, (iv) ssDNA binding with multiple to multiple binding sites on RAD52 monomers, oligomers, rings or larger aggregates, and (v) hybridization with and without bound protein. Unknown parameters include the affinity of the two binding sites of short RAD52 oligomers to ssDNA and their ends. Affinities and cooperativity likely differ depending on the RAD52 oligomeric state and DNA substrate^37,67^. Also, RAD52 may undergo conformational changes upon oligomerization and/or DNA binding similar to its homologue Red*β* ^37^. Since there are many competing biochemical reactions, most of which are not quantitatively characterized, annealing, homology search, and inhibition reactions are still challenging to understand. Yet, the extensive literature and our findings narrow down possible explanations at physiological conditions.

### RAD52-ring-DNA biochemistry

Our data does not strongly support a ring-based annealing mechanism. Multiple results rather indicate that rings inhibit annealing by immobilizing ssDNA on them. By this immobilization, bases may be made inaccessible for homology search especially for RAD52(209) having a higher propensity to form rings. Our data show that two or more ssDNA can stably bind to rings. All ring-based annealing models assume that only one strand is bound to one ring and the other strand is either bound to another ring or may bind to the second binding site. However, if both sites are already occupied by two strands of the same type after incubation with only one of the strands in our *in vitro* experiments (Fig. 7–9), such mechanisms are impossible to explain our *in vitro* data. Also, there is steric hindrance between two rings keeping DNA strands too far apart for homology detection^27^. Homology detection is also difficult due to the ring curvature. Furthermore, if two identical strands are wrapped around rings, one of them should also be bound to the outer site. There, the ssDNA is twisted around itself being inaccessible for base pairing^27^. We found that longer ssDNA strands that may wrap around rings were bound stably (Fig. 6). More stable binding to rings compared to short oligomers is expected based on the increased number of binding sites and the data for RAD52’s homologue^37^. Thus, dissociation or partial peeling off^35^ from rings to circumvent steric or curvature issues during annealing is unlikely. Therefore, in particular for high protein concentrations that favor ring formation, a ring-based inhibition scenario is likely independent of whether one or both strands were incubated with the protein prior to the annealing reaction.

Consistently, reduced product yield correlated with high, ring-promoting concentrations (Fig. 5 and previous reports^9,35,51,52^) supporting a ring-inhibition mechanism. Additional aggregation of rings, which we also observed for high RAD52 concentrations (Fig. 7 and S5), has been suggested to be the reason for the decrease of product. If rings promote annealing, product yield should be proportional to ring concentration. At concentrations between 10–100 nM, we did not yet observe much aggregation (less than 0.5 % of protomers were in aggregates). Over this concentration range, the ring concentration increased by 1800 % (Fig. 3b) that should result in a similar increase in product yield below the *K*_*d*_ of the reaction (≈ 100 nM for the first step of ssDNA binding Fig. 4a). However, product yield increased only by about 25 % (Fig. 5c). Thus, the law of mass action would be violated for a ring-based annealing mechanism. By contrast, the RAD52 short oligomer concentrations remained constant over this range, with a 25 % increase being possible within error bars. Apart from a product yield increase with ring concentration, ring-based annealing should also result in a higher cooperativity of ssDNA binding that is inconsistent with our data (Fig. 4).

For high protein concentrations, ring-mediated annealing inhibition seems the most plausible explanation. At our lowest protein concentration tested, there were not enough rings present to inhibit annealing when both strands were incubated with the protein prior to mixing, in particular not for the full-length protein. For example, at 10 nM RAD52, the ring concentration was only about 0.5 nM (Fig. 3b) compared to the 10 nM concentration of the ssDNA. Thus, under these conditions, there was even an excess of ssDNA that might disassemble rings (Fig. 9). Since the concentration of RAD52 rings was much smaller than that of the ssDNA, there were also not enough rings present to promote product yield by a factor two (Fig. 5c). Recycling of rings after a successful annealing reaction for multiple turnovers of annealing is also not likely since (i) DNA was stably bound to rings, and (ii) once DNA is dissociated form a ring, the situation is equivalent to mixing all components directly, for which we did not observe an increase in product yield (Fig. 5d). Thus, at 10 nM RAD52, rings likely neither significantly promoted nor inhibited annealing. Therefore, a second inhibition mechanism must exist.

For inhibition at low protein concentrations, we favor the idea that end-bound monomers and/or dimers block annealing. This hypothesis is supported by the end-binding preference of RAD52^28,65^ and our measurement that RAD52 monomers and dimers are the only short oligomers present at low concentrations (Fig. 3). If complementary ends of both strands have RAD52 bound, annealing may be inhibited either due to steric hindrance with two RAD52-bound oligonucleotides being too far apart when bound to the inner groove or not being available for pairing when bound to the outer site because of the twisted nature of the bound ssDNA^27^. If only one strand has RAD52 bound, a RAD52-free complementary strand may scan the nucleoprotein filament for homology and anneal if the filament is sufficiently long to ensure uniqueness. Such a scenario would imply that the promoting of the annealing reaction is mainly due to enhanced hybridization initiated at the ends of the strands. Annealing from the ends would match the *in vivo* situation. In summary, the above arguments strongly suggest that annealing is promoted by RAD52 monomers or short oligomers instead of rings.

### A dynamic multistep, short-oligomer-mediated annealing mechanism

Based on our data and with reference to the Red*β* model^26,37,46,47^, we propose the following annealing and homology search model for RAD52 (Fig. 10). The model is based on sequential association of RAD52 monomers with each other and with DNA resulting in a short nucleoprotein filament that enables homology search with a complementary strand free of RAD52. Each step corresponds to a unique complex or binding situation that allow annealing and mismatch discrimination without ATP. Annealing occurs in multiple phases or steps to ensure that unique homology is detected. Both rings and larger aggregates of rings with ssDNA inhibit annealing (Fig. 10).

**Figure 10.**
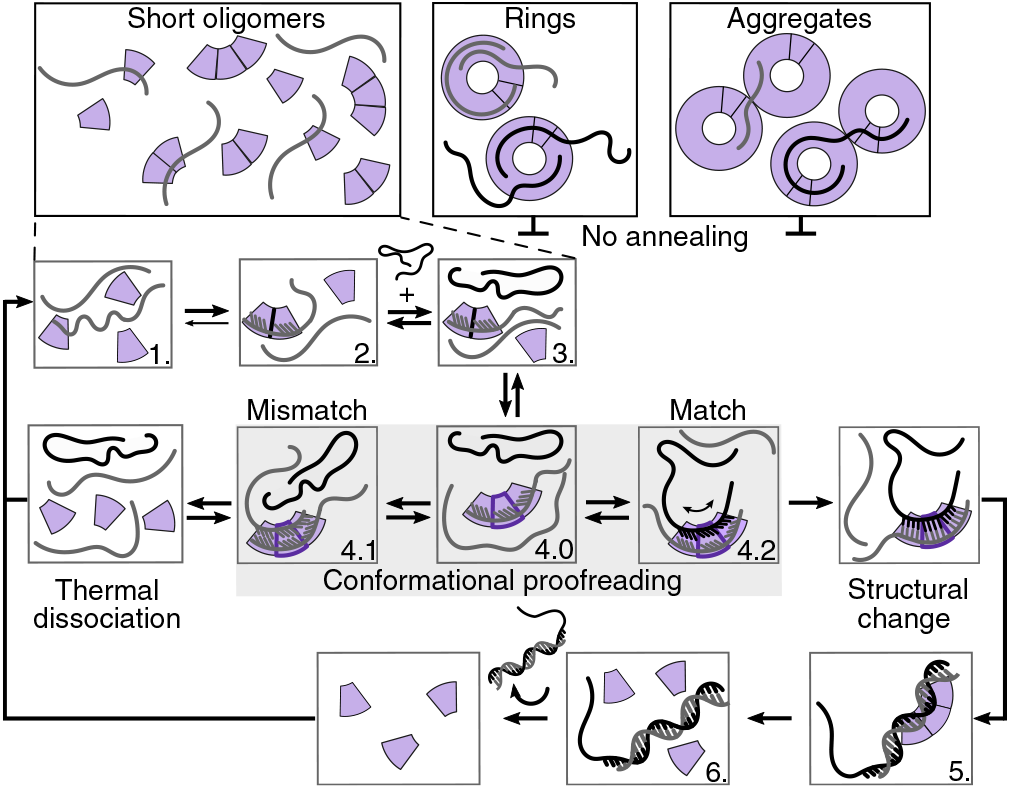
RAD52 single-strand annealing mechanism. Short RAD52 oligomers (purple annulus sectors) anneal complementary ssDNA strands (gray and black lines) in a multi-step, catalytic mechanism. Rings or their aggregates inhibit annealing. See text for details.

To promote annealing, in the first step, monomers bind ssDNA with a preference to DNA ends (Step 1.). *In vivo* it can replace RPA, potentially recruit RAD51, and protect ends against exonucleases^12,28,65,69^. Since ssDNA binding to the inner binding site of RAD52 is endothermic and to the outer exothermic with a higher affinity^27^, we propose that ssDNA is first bound to the outer binding site. At this site, ssDNA is bound in a helical fashion^27^ and may not be accessible for base pairing. In this manner, RAD52 monomers bound to ssDNA inhibit annealing for a lack of sufficient homology^39^ and also protect ssDNA from degradation. Also when multiple non-consecutive monomers are bound to complementary strands, annealing is inhibited consistent with our data on incubating both strands with the protein prior to mixing them (Fig. 5).

In the second step, another RAD52 monomer binds in a consecutive manner to a RAD52-monomer-bound ssDNA forming a RAD52-dimer-bound ssDNA (Step 2.). Since DNA binding to the inner site of RAD52 is endothermic, we propose that the energetic gain in self-association of the two RAD52 monomers is used to transfer the ssDNA from the outer to the inner binding site. We think that dimer formation and ssDNA transfer to the inner site is a coupled process supported by our finding that ssDNA-binding is cooperative (Fig. 4). The temperature dependence of cooperativity (Fig. S3 and Table S3) suggests that the transfer involves the transition over a significant activation barrier. One reason for the endothermic binding to the inner site may be that the ssDNA is compressed to a near B-form structure^27^ enabling conformational proof-reading^37,70–72^ with a complementary strand. The coupled process and transition barrier ensure that dimer formation occurs on the filament and not through a RAD52 oligomer directly binding to the ssDNA from solution. This mechanism ensures consecutive binding and formation of a filament without gaps and may make the binding of larger oligomers unfavorable. Larger oligomers, in particular when bound in a non-consecutive, off-register fashion may be difficult to dissociate after homology detection.

The RAD52-dimer-bound ssDNA provides eight compressed bases for homology search. While eight bases are an efficient unit for search, it is not yet sufficiently long for uniqueness^39^. Also, hybridization of eight bases is not yet thermally stable. The gain in free energy due to hybridization of a ssDNA with a compressed, near-B-form, RAD52-bound ssDNA is likely even smaller resulting in an even less stable complex. Thus, only complementary strands may bind at all for a sufficiently long homology search time. In this manner, such a conformational proofreading discriminates mismatches. A weaker binding may also allow for a complementary strand to scan the RAD52-bound strand, for example, via one-dimensional diffusion. That only one strand but not its complementary strand has RAD52 bound is supported by our incubation-order experiments (Fig. 5). Since binding of a complementary strand to the RAD52-dimer-bound ssDNA is likely energetically less favorable compared to the binding to bare ssDNA, annealing is initially not accelerated or promoted but slowed down. Such an effect was observed for the RAD52 phage homologue Red*β* ^37^ and is a hallmark for conformational proofreading. The RAD52-dimer-bound ssDNA is a unique structure in the sense that both RAD52 protomers only have one lateral protein-protein contact. Yet, this complex is not sufficient for identifying a unique sequence of homology.

In a third step, at least one more RAD52 monomer binds to the RAD52-dimer-bound ssDNA, again in a consecutive, cooperative manner forming a short nucleoprotein filament containing three RAD52 protomers (Step 3.). Such a nucleoprotein filament is now long enough for unique homology identification^39^. With three or more protomers bound, the complex is again special in the sense that now at least one protomer in the middle of the nucleoprotein filament has two protein-protein contacts. This middle RAD52 protomer should have a conformation similar to the one of the protomers in a ring while the RAD52 protomers at the two ends of the short nucleoprotein filament may have a different conformation. Indirect evidence for different conformations or a different strength of the inter-protein interaction stem from experiments that disrupted rings by drugs resulting surprisingly in dimers and not monomers^30^. In general, for filament growth the dimer association constant is often different from the association constant of subsequent monomers and was also observed for Red*β* ^37, 64^.

Once a three protomer containing nucleoprotein filament has formed, a second strand without any bound RAD52 can diffuse along the nucleoprotein filament in search for homology (Step 4.). In case of a mismatch and invoking conformational proofreading, base pairing energies are not sufficient to form a thermally stable dsDNA (Step 4.1). Thus, strands with a mismatch are rejected by thermal fluctuations. This is an essential step in homology detection for a protein that is not an ATPase. If strands are complementary, base-pairing energies are sufficient for a stable complex. In a subsequent step, we propose that the annealing energy is used for a structural change of RAD52 (Step 4.2). Such a change upon filament formation was found for Red*β* by circular dichroism measurements and may be the reason for the remarkable stability of Red*β* nucleoprotein filaments after homology is detected^37^. For human RAD52, we propose that such a structural change only occurs for short nucleoprotein filaments and in such a manner that the dsDNA product is not clamped by RAD52 but released from it (Step 5. and 6.). Such a release would be consistent with our annealing gel measurements up to 200 nM of protein for which rings were not yet dominating and for which we observed an increase in free product yield (Fig. 6). Weak interactions of dsDNA with RAD52 are also consistent with previous work^32^. Thus, in a multistep process, the step-wise assembly of a nucleoprotein filament enables to harness both self-association and base-pairing energies for an efficient, robust homology detection and release of a successfully annealed dsDNA product. In this sense, RAD52 may truly act as a catalyst to promote annealing.

### Concluding remarks

While we cannot directly visualize or detect the annealing process by RAD52 monomers or short oligomers, our data is most consistent with such a model and inconsistent with a ring-mediated annealing model. The C-terminal domain of RAD52 modulates the self-association of RAD52 resulting in more monomers and less rings of the full-length protein compared to the truncated one. According to our model, both factors promote annealing which is supported by our data (Fig. 5c). *In vivo* the C-terminal domain is key for interacting with RPA and RAD51. It may also interact with the complementary strand mediating an efficient scanning process for homology detection and/or the release of the dsDNA product. For successful annealing, both DNA base pairing and RAD52 self-association is important. If the latter is disrupted, the protein does—according to our model—not promote annealing anymore, which is consistent with experiments using small-molecule inhibitors^14,30^. Estimated physiological concentrations are in the lower nanomolar range. At these concentrations, we observed only few larger oligomers and rings. These may be necessary for localization to the nucleus. Rings may also act as a protein reservoir that is disassembled upon DNA damage.

To test our RAD52 short-oligomer-mediated model, further experiments are necessary ideally visualizing the process for example by simultaneous mass photometry and single-molecule fluorescent imaging, ideally also in the presence of RPA. We believe that our current findings contribute to a better understanding of the single-strand annealing mechanism by RAD52 and will help to develop drugs that efficiently target it for cancer treatments.

## Code availability

Python script for the mass photometry data analysis and statistics: https://github.com/tobiasjj/iscam_analysis

## Data availability

The authors confirm that the data supporting the findings of this study are available within the article and its supplementary materials.

## Acknowledgements

We thank the group of Dr. John Weir for letting us use the OneMP setup. We thank Anita Jannasch, Carolina Carrasco Pulido, Shu Yao Leong, Yannic Lurz, Serapion Pyrpassopoulos, and Hauke Drechsler for comments on the manuscript. M.S.K. and P.K. acknowledge funding from the European Research Council (Consolidator Grant PHOTOMASS 819593) and an Engineering and Physical Sciences Research Council Leadership Fellowship (EP/T03419X/1). For the purpose of Open Access, the author has applied a CC BY public copyright licence to any Author Accepted Manuscript (AAM) version arising from this submission. M.A.K. acknowledges financial support from the International Max Planck Research Schools from Molecules to Organisms, Max Planck Institute for Developmental Biology, Tübingen and from the University of Tübingen.

## Author contributions statement

The project was designed by E.S. and P.K. Preliminary mass photometry measurements with RAD52(209) and the followup analysis were performed by T.J.J. and M.S.K. Mass photometry measurements on RAD52(209) and RAD52 were performed by M.A.K. Biochemical assays were designed and performed by M.A.K. The Python script for the mass photometry data analysis and statistics was written by T.J.J. and M.A.K. Proteins were purified by S.S. Data analysis of mass photometry and biochemical assays was performed by M.A.K. and E.S. Figures were prepared by M.A.K and E.S. The manuscript was written and edited by M.A.K., A.F.S. and E.S. with comments from all authors.

## Competing interests

None declared.

## Notes

### Competing Interest Statement

The authors have declared no competing interest.

