## Supporting Information for "Short oligomers rather than rings of human RAD52 promote single-strand annealing"

### Supplementary Information

**Table S1.** List of oligonucleotides.

| Name | Sequence |
|---|---|
| 32 nt 3'-ATTO565 | GCTCTAAGCCATCCGCAAAAATGACCTCTTAT |
| 48 nt 3'-ALEXA488 | GCAATTAAGCTCTAAGCCATCCGCAAAAATGACCTCTTATCAAAAGGA |
| 60 nt 3'-ATTO680 | TCCTTTTGATAAGAGGTCATTTTTGCGGATGGCTTAGAGCTTAATTGCTTTTTTTTTTTT |
| 70 nt 3'-ATTO565 | TTTAATAATTTACTTTATTTTCTATGTCTATTCATTTACTTATTGTGTATTATCCTTATACTTACTTAC |
| 83 nt[a] | TTTATATAATTTACTTTATTTTCTATGTCTATTCATTTACTTATTGTGTATTATCCTTATACTTACTTAC TTTATGTTTATTT |

[a] Mazina *et al.*, Ref. 6 in main text.

**Table S2.** Molecular weights for the RAD52 protomers. The table contains estimated weights based on the protein sequence and the values obtained by mass photometry.

| Oligomer (number of protomers) | Sequence weight (kDa) | Measured weight (kDa) |
|---|---|---|
| Monomer (1) | 48.4 | $43 \pm 2$ |
| Dimer (2) | 96.8 | $89 \pm 3$ |
| Trimer (3) | 145.2 | $140 \pm 5$ |
| Tetramer (4) | 193.6 | $185 \pm 4$ |
| Nonamer (9) | 435.6 | $447 \pm 3$ |
| Decamer (10) | 484 | $491 \pm 2$ |
| Undecamer (11) | 532.4 | $532 \pm 2$ |
| Dodecamer (12) | 580.9 | $573 \pm 5$ |

**Table S3.** Hill equation fit parameters for the single-stranded DNA binding reactions. Binding affinities of RAD52(209) and RAD52 were measured for different lengths of oligonucleotides (32, 60, and 70 nt) at different temperatures (25 and 37 °C). Each condition was measured at least 3 times. Mean values with SEM are plotted as the function of RAD52 concentration (Fig. S3). Experimental data was fitted to Hill and Michaelis-Menten equations, fits were weighted based on SEM, $V_{max}$ was fixed to 100 %.

| | RAD52(209) | | RAD52 | |
|---|---|---|---|---|
| Number nt ($T$ in °C) | $K_d$ | $H$ | $K_d$ | $H$ |
| 32 nt (25) | $169 \pm 24$ | $1.5 \pm 0.2$ | $115 \pm 22$ | $1.4 \pm 0.2$ |
| 70 nt (25) | $109 \pm 23$ | $1.6 \pm 0.3$ | $53 \pm 8$ | - |
| 32 nt (37) | $246 \pm 33$ | $3.1 \pm 0.5$ | $135 \pm 4$ | $2.7 \pm 0.2$ |
| 60 nt (37) | $127 \pm 10$ | $4.1 \pm 0.4$ | $112.4 \pm 0.1$ | $3.3 \pm 0.2$ |

**Table S4.** Fit parameters for second-order reaction kinetics $2Akt/(1/A+2kt)$ in Fig. 4c

| Reaction | Amplitude $A$ | Rate constant $k$ (min$^{-1}$) |
|---|---|---|
| Control | $3.3 \pm 0.2$ | $0.068 \pm 0.009$ |
| RAD52(209) | $3.59 \pm 0.06$ | $0.39 \pm 0.08$ |
| RAD52 | $5.16 \pm 0.03$ | $2.0 \pm 0.3$ |

**Table S5.** Shifts of molecular weights of RAD52(209) single ring peak after the binding of single-stranded DNA of different lengths and during the annealing reaction. The protein concentration was 100 nM, ssDNA concentrations were 10 nM. The table contains mean values with SEM after several repeats (32 nt - 7, 48 nt - 6, 60 nt - 6, 83 nt - 6, annealing reaction - 9).

| Length in nt (kDa) | MW shift (kDa) |
|---|---|
| 32 (9.8) | $20 \pm 1$ |
| 48 (14.8) | $32 \pm 2$ |
| 60 (18.6) | $42 \pm 1$ |
| 83 (25.4) | $40 \pm 4$ |
| 60 (preincubated) + 32 (28.4) | $42 \pm 2$ |

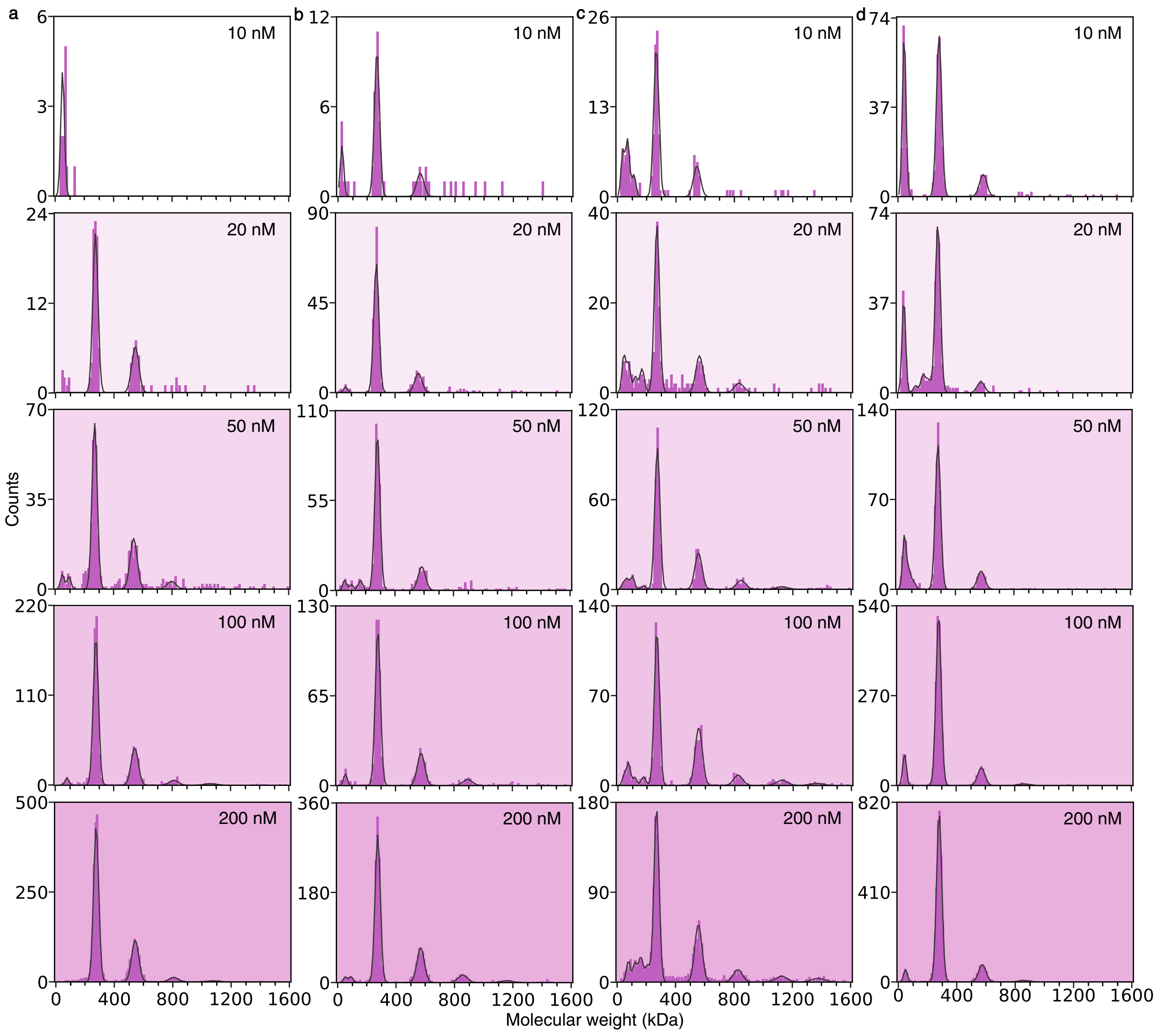


**Figure S1.** Oligomeric states of RAD52(209) at different concentrations (10, 20, 50, 100 and 200 nM) on different days (**a**)-(**d**). Each plot is a cumulative distribution from 2–3 measurements from the same day.

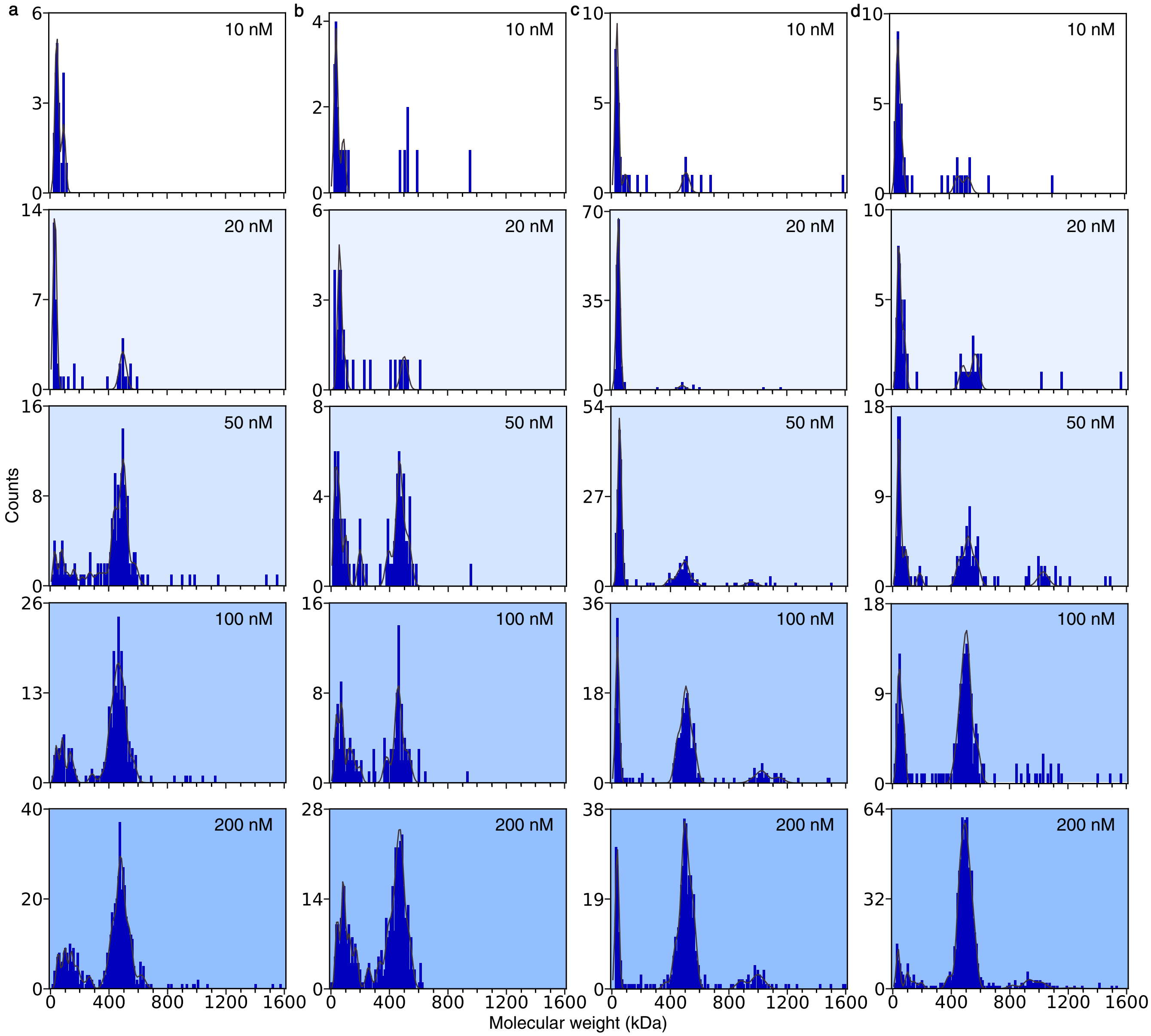


**Figure S2.** Oligomeric states of RAD52 at different concentrations (10, 20, 50, 100 and 200 nM) on different days **(a)**- **(d)**. Each plot is a cumulative distribution from 2–3 measurements from the same day.

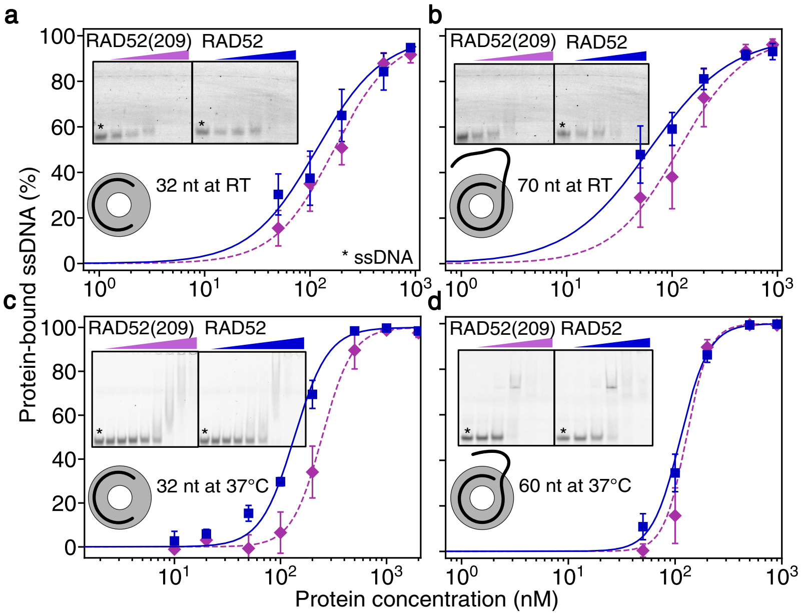


**Figure S3.** Electrophoretic mobility shift assays of ssDNA binding to RAD52(209) and RAD52. Three different lengths were tested, 32, 60, and 70 nt, which correspond to 8, 15, and 17.5 binding sites, respectively. DNA binding was tested at 25 °C with 32 nt (5 repeats) **(a)** and 70 nt (4 repeats) **(b)**, and at 37 °C with 32 nt (4 repeats) **(c)** and 60 nt (3 repeats) **(d)**. See Table S2 for Hill equation fit parameters.

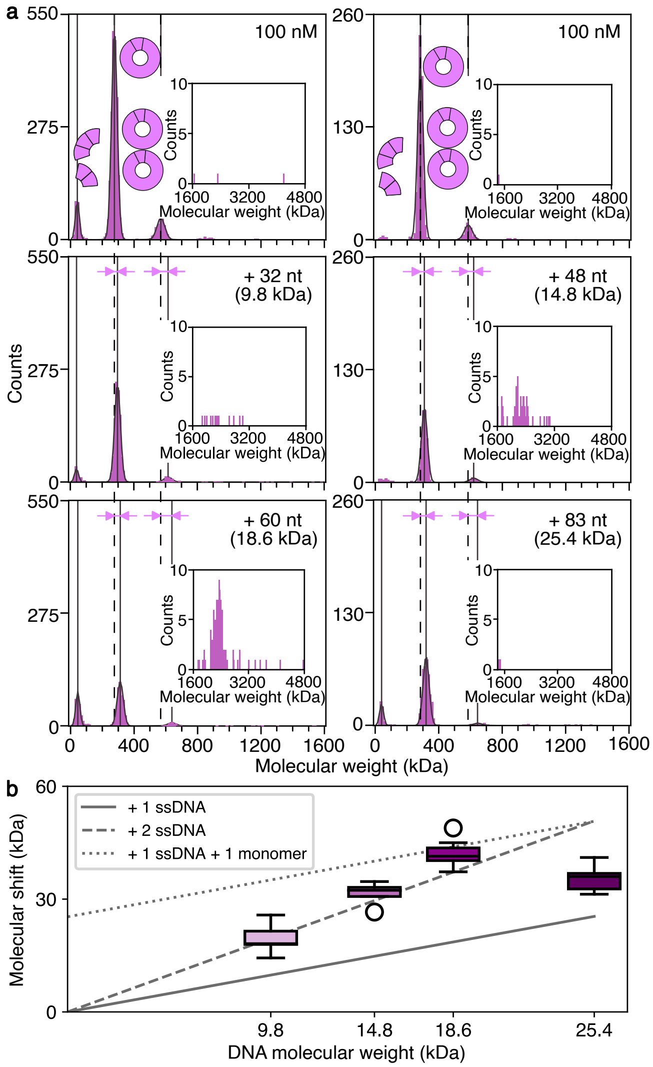


**Figure S4.** MP data presents RAD52(209) binding to single-stranded DNA molecules with different lengths: 32 (9.8 kDa), 48 (14.8 kDa), 60 (18.6 kDa), and 83 nt (25.4 kDa)**(a)**. Corresponding MW shifts marked with pink arrows and quantified **(b)**. The plot contains MW shifts mean values with SEM from different repeats. RAD52(209) concentration was 100 nM. DNA concentrations of were 10 nM.

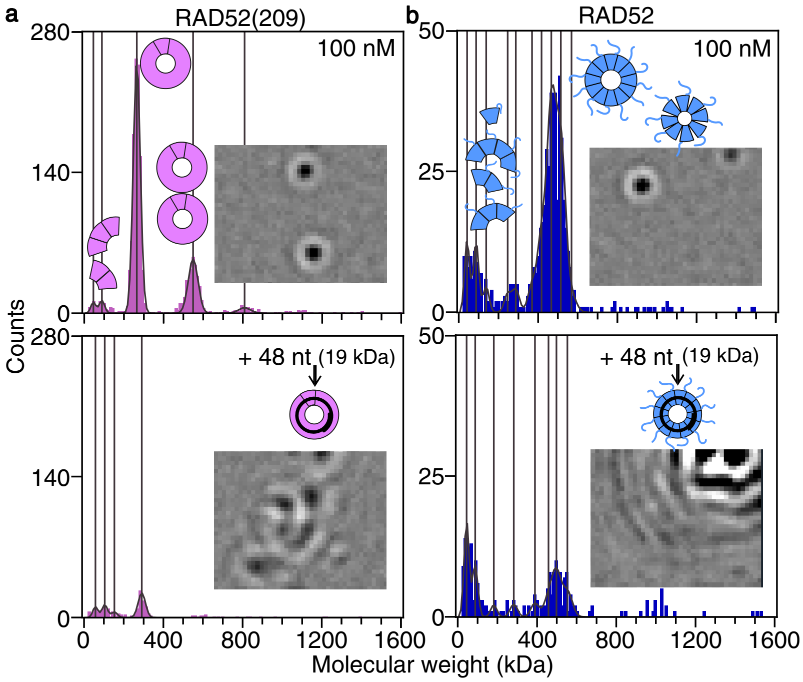


**Figure S5.** Mass photometry data of RAD52(209) **(a)** and RAD52 **(b)** at 100 nM and with ssDNA bound (48 nt). Formation of large clusters was observed (insert). DNA concentration was 10 nM. Each dataset is a cumulative distribution from three repeats.
